# Interpretable Multiomics Machine Learning Identifies GSDMB-Associated Epigenetic Repression and Reduced Immune Activity in Metastatic Colorectal Cancer

**DOI:** 10.64898/2026.09.01.748124

**Authors:** Matheus da Silveira Costa, Henrique Izaias Marcelo, Gabriel Albanese Kafouri, Vinicius de Camargo

## Abstract

**Background:** Colorectal cancer (CRC) is a major cause of cancer-related mortality, with distant metastasis strongly associated with poor clinical outcomes. Integrating transcriptomic and epigenomic data through machine learning may improve the molecular characterization of metastatic CRC.

**Methods:** We analyzed 518 primary tumors from the TCGA-COAD/READ cohort (436 non-metastatic [M0] and 82 metastatic [M1]) with matched RNA-seq and DNA methylation data. Five machine learning classifiers were evaluated for discrimination of metastatic status using stratified nested cross-validation. Model explainability was assessed using model coefficients for linear classifiers and SHAP-based feature importance for tree-based models. Differential expression, functional enrichment, methylation-expression correlation, immune-related transcriptional scoring, statistical mediation, and survival analyses were subsequently performed.

**Results:** Integrated RNA-seq and DNA methylation showed the strongest discrimination between M0 and M1 tumors, with SVM reaching a ROC-AUC of 0.787 +/− 0.047. Six features - ARC, ASPDH, C13orf15, C4orf23, GPATCH3, and cg12040555 - ranked among the top 20 predictors across four models with consistent directions. cg10057218 methylation was inversely correlated with GSDMB expression (Spearman rho = −0.589) and increased in M1 tumors. Statistical mediation indicated a significant indirect association between M1 status and reduced GSDMB expression through cg10057218 methylation (indirect effect = −0.370; 95% CI [−0.519, −0.224]; proportion mediated = 82.1%). M1 tumors also exhibited reduced immune-related transcriptional activity. A 20-feature canonical molecular score separated M1 from M0 tumors (in-sample ROC-AUC = 0.918), while the corresponding Logistic Regression model achieved a nested cross-validated ROC-AUC of 0.781 +/− 0.046. Higher scores were associated with shorter overall survival (log-rank p = 1.67 x 10^-7).

**Conclusions:** Transcriptomic and DNA methylation integration identified a consistently prioritized multiomics feature set associated with metastatic CRC. The findings highlight coordinated molecular and immune-related differences between M0 and M1 tumors and identify cg10057218-associated GSDMB repression as a candidate epigenetic feature of metastatic disease.

## 1. Introduction

Colorectal cancer (CRC) is one of the most frequently diagnosed malignancies worldwide and remains a leading cause of cancer-related mortality. In 2022, CRC accounted for approximately 1.93 million new cases and more than 904,000 deaths globally [1,2]. Although localized CRC is often successfully treated, prognosis deteriorates markedly after distant dissemination, with five-year relative survival decreasing from more than 90% for localized disease to approximately 15% for metastatic disease [3,4]. Understanding the molecular features associated with distant metastasis therefore remains an important challenge in CRC research.

CRC is molecularly heterogeneous, encompassing distinct patterns of genomic alteration, chromosomal and microsatellite instability, DNA methylation, transcriptional activity, and tumor–immune interactions [5,6]. Large-scale initiatives such as The Cancer Genome Atlas (TCGA) have enabled these molecular dimensions to be characterized within the same tumors [5]. However, metastatic competence arises from coordinated alterations across multiple biological processes, including cellular plasticity, angiogenesis, immune evasion, metabolic adaptation, and resistance to cell death [7–10]. Consequently, individual omics layers may capture only part of the molecular variation associated with metastatic disease.

Among these regulatory layers, DNA methylation is particularly relevant because it can contribute to stable changes in gene expression without altering the underlying DNA sequence [11,12]. Aberrant DNA methylation is a well-established feature of colorectal tumorigenesis, with effects on transcriptional regulation, genomic stability, and tumor– immune interactions [5,11–13]. Integrating methylation with transcriptomic data may therefore help identify epigenetic alterations that are not only associated with metastatic status but also linked to functional changes in gene expression.

One potentially relevant biological context involves programmed and inflammatory cell death. Gasdermins are pore-forming proteins involved in inflammatory cell-death pathways and have context-dependent roles in tumor biology, immune regulation, and therapeutic response [14–18]. Among them, gasdermin B (GSDMB) has been implicated in cancer progression and regulation of cell-death-related processes [17,18]. However, the relationship between GSDMB expression, DNA methylation, and metastatic CRC remains insufficiently characterized. More broadly, whether methylation-associated transcriptional alterations in metastatic CRC occur alongside changes in immune-related transcriptional programs remains an open question.

Machine learning provides a framework for integrating high-dimensional molecular data and identifying multivariable patterns associated with disease phenotypes. However, predictive performance alone provides limited biological insight. Explainable machine-learning approaches, including model coefficients and SHAP-based feature importance, can identify the molecular variables contributing most strongly to classification and enable prioritization of features for subsequent biological investigation [19]. Combining such approaches with methylation–expression analysis, functional enrichment, and immune-related transcriptional profiling may therefore bridge predictive modeling and biological interpretation.

In this study, we investigated whether integration of RNA-seq and DNA methylation data improves discrimination between non-metastatic (M0) and metastatic (M1) primary CRC tumors and enables identification of complementary molecular features associated with metastatic status. Using TCGA-COAD/READ, we compared five machine-learning classifiers under nested cross-validation and applied model-specific explainability approaches to identify robust predictive features. We subsequently integrated methylation–expression correlation, differential expression, functional enrichment, immune-related transcriptional scoring, statistical mediation, and survival analyses to characterize the biological context of the identified signature, with particular emphasis on the relationship between GSDMB-associated methylation and gene expression.

## 2. Methods

### 2.1. Data Source and Cohort

TCGA-COAD/READ data were obtained from cBioPortal [24]. The cohort comprised 523 patients with available RNA-seq data and defined pathological metastatic status, of whom 518 also had available DNA methylation profiles. Metastatic status was defined according to the PATH_M_STAGE variable. Patients classified as M1, M1a, or M1b were grouped as metastatic (M1), whereas those classified as M0 were grouped as non-metastatic (M0). Patients with MX classification were excluded. Importantly, the molecular analyses were restricted to primary colorectal tumor samples; no samples obtained from metastatic lesions were included. Thus, the M1 group represents primary tumors from patients classified as having distant metastatic disease, rather than tissue collected from metastatic sites.

Only primary tumor samples, identified by the TCGA sample-type code 01, were retained. Patient identifiers were derived from TCGA sample barcodes, and multiple primary-tumor samples mapping to the same patient, if present, were averaged at the patient level. In the dataset analyzed here, no duplicate primary-tumor samples were present after filtering.

### 2.2. Data Preprocessing

RNA-seq data, provided as RSEM-normalized expression values, were log₂-transformed as log₂(RSEM + 1). DNA methylation data, derived from merged HM27/HM450K platforms, were provided as beta values ranging from 0 to 1. RPPA data were represented as patient-level protein abundance measurements, retaining primary tumor samples and averaging duplicate samples mapped to the same patient. Somatic mutation data were represented as a binary patient-by-gene matrix, in which a value of 1 indicated the presence of at least one somatic mutation in a given gene and 0 indicated its absence; genes mutated in fewer than 3% of patients were excluded.

For machine learning analyses, median imputation and feature standardization were performed within the cross-validation framework for all omics configurations. For RNA-seq, DNA methylation, and the integrated RNA-seq + methylation configuration, zero-variance features were removed and univariate feature selection was subsequently performed using the ANOVA F-statistic. The number of selected features was 200 for RNA-seq, 100 for DNA methylation, and 300 for the integrated RNA-seq + methylation model. No additional SelectKBest feature-selection step was applied to RPPA or somatic mutation data. All preprocessing steps requiring parameter estimation were fitted exclusively on the training folds and then applied to the corresponding validation folds to prevent data leakage.

### 2.3. Machine Learning Models and Cross-Validation

Five classifiers were evaluated: Logistic Regression, linear-kernel Support Vector Machine (SVM), LightGBM, XGBoost, and Random Forest. Class imbalance was addressed using class_weight = “balanced” for Logistic Regression, SVM, LightGBM, and Random Forest. For XGBoost, scale_pos_weight was calculated separately within each outer training fold as the ratio of M0 to M1 samples, thereby preventing information from the held-out outer test fold from contributing to class-weight specification.

Model performance was evaluated using nested stratified cross-validation, with a 5-fold outer loop for performance estimation and a 3-fold inner loop for hyperparameter optimization. Within each outer training fold, hyperparameters were optimized using RandomizedSearchCV with 30 iterations and ROC-AUC as the optimization metric. The optimized model was then evaluated on the corresponding held-out outer fold. Performance metrics included ROC-AUC, PR-AUC, Brier score, sensitivity, specificity, Youden index, precision for the M1 class, F1 score for the metastatic class, and balanced accuracy. For threshold-dependent metrics, the Youden-optimal threshold was determined within each outer training fold using out-of-fold predictions generated by three-fold stratified cross-validation of the tuned pipeline. The resulting fold-specific threshold was then applied to the corresponding outer-test probabilities to compute sensitivity, specificity, precision, F1 score, balanced accuracy, and Youden index. The outer test fold was not used for threshold selection.

### 2.4. Explainability Analysis

Random Forest was included in the comparative performance analysis but was not carried forward to the feature-level consensus analysis because it showed lower discriminative performance in the integrated RNA-seq + DNA methylation setting. Therefore, feature importance and consensus analyses were restricted to logistic regression, linear SVM, LightGBM, and XGBoost, which showed superior or more competitive ROC-AUC performance.

For tree-based models, including LightGBM and XGBoost, SHAP values were computed using TreeExplainer on models trained on the full dataset. Feature importance was quantified as the mean absolute SHAP value, defined as mean(|SHAP|). Feature direction was determined by the Spearman correlation between feature values and their corresponding SHAP values across all samples, following the coloring convention used in SHAP beeswarm plots.

For logistic regression with L2 regularization and linear-kernel support vector machine, feature importance was defined as the absolute value of the standardized model coefficient, whereas the coefficient sign was used to indicate the direction of association.

To identify consistently prioritized predictive features, the top 20 features from each of the four models were compared. Features appearing among the top 20 predictors in all four models were defined as cross-model consensus features.

### 2.5. Differential Expression Analysis

Differential gene expression between metastatic (M1) and non-metastatic (M0) tumors was assessed using RNA-seq data from the matched cohort. For each gene, expression values were compared between M1 and M0 tumors using a two-sided Mann– Whitney U test. Genes with fewer than five valid observations in either group were excluded. Expression differences were quantified as the difference between median log₂-transformed expression values in M1 and M0 tumors (Δlog₂ expression = median M1 − median M0). P-values were adjusted for multiple testing using the Benjamini–Hochberg false discovery rate (FDR) procedure, and genes with FDR < 0.05 were considered differentially expressed. Among significant genes, positive Δlog₂ expression values indicated higher expression in M1 tumors, whereas negative values indicated lower expression in M1 tumors. For subsequent exploratory functional enrichment analyses, broader directional gene sets were defined using FDR < 0.10 and Δlog₂ expression > 0 for M1-upregulated genes or Δlog₂ expression < 0 for M1-downregulated genes.

### 2.6. Functional Enrichment Analysis

Over-representation analysis was performed using gseapy.enrichr. Functional enrichment of prioritized M1-associated RNA-seq features derived from the machine learning models was evaluated against MSigDB Hallmark 2020, Gene Ontology Biological Process 2023, KEGG 2021 Human, and Reactome 2022 gene-set collections. For this model-derived analysis, the enrichment background was restricted to the 199 RNA-seq genes retained by the integrated modeling pipeline.

In a complementary analysis, functional enrichment of genes showing directional expression differences between M1 and M0 tumors was evaluated using the broader gene sets defined by FDR < 0.10 and Δlog₂ expression > 0 or < 0 for M1-upregulated and M1-downregulated genes, respectively. These gene sets were analyzed against Gene Ontology Biological Process 2023, Gene Ontology Molecular Function 2023, KEGG 2021 Human, Reactome 2022, and MSigDB Hallmark 2020. Enrichment results with Benjamini– Hochberg-adjusted p-values < 0.05 were considered statistically significant.

### 2.7. Methylation–Expression Correlation Analysis

For CpG probes annotated to a single gene based on the NAME field of the HM27/HM450K methylation annotation, methylation–expression correlation analysis was performed when the corresponding gene was also present in the RNA-seq dataset. Probes annotated to multiple genes were excluded from this analysis. Spearman’s correlation coefficient was calculated between CpG methylation β-values and log₂(RSEM + 1)-transformed gene expression values across the matched multiomics cohort. Only CpG– gene pairs with at least 30 valid paired observations were retained. In total, 19,150 CpG– gene pairs corresponding to 11,245 unique genes were analyzed. P-values from the full set of 19,150 correlations were adjusted for multiple testing using the Benjamini–Hochberg false discovery rate procedure.

To prioritize methylation features identified by the machine-learning analyses, the 101 CpG probes retained within the common 300-feature set across the four integrated models were intersected with the methylation–expression correlation results. Of these, 86 had eligible CpG–gene pairs under the single-gene annotation criterion. As a complementary assessment of feature robustness, cross-validation feature-importance stability was evaluated using three classifiers (LightGBM, XGBoost, and logistic regression) across five stratified cross-validation folds. Within each fold, preprocessing and feature selection were performed using the training data before model fitting. For each fold–model combination, the 50 features with the highest model-specific absolute importance were retained, yielding 15 feature-ranking opportunities (5 folds × 3 models). For each feature, frequency was defined as the number of fold–model combinations in which it appeared among the top 50 features, and stability was calculated as frequency divided by 15. Mean rank and mean importance were calculated across the fold–model combinations in which the feature was retained.

Candidate CpG–gene pairs were subsequently evaluated by integrating complementary evidence from model-based feature prioritization, cross-validation feature-importance stability, methylation–expression correlation magnitude, and multiple-testing-adjusted statistical significance. Strong inverse methylation–expression relationships were prioritized as candidates for further investigation.

### 2.8. Immune-Related Transcriptional Score

To assess immune-related transcriptional activity, a curated 29-gene panel was assembled based on established biological functions and supported by previously published tumor immune-expression signatures [20, 21]. The genes were grouped into four functional modules: cytotoxic T/NK activity (CD8A, CD8B, GZMB, PRF1, NKG7, GNLY, GZMA, GZMK) [20,21], interferon-γ response (IFNG, IRF1, CXCL9, CXCL10, CXCL11, STAT1, IDO1) [20], antigen presentation (HLA-A, HLA-B, HLA-C, HLA-DRA, HLA-DRB1, B2M, CD74) [20,21], and immune checkpoint/Treg-associated expression (PDCD1, CD274, CTLA4, FOXP3, TIGIT, LAG3, HAVCR2) [20,21]. Gene expression values were standardized within the matched 518-patient multiomics cohort by calculating gene-level z-scores across patients. For each patient, module-specific scores were calculated as the mean z-score of the genes assigned to each functional category, and the composite immune-related transcriptional score was calculated as the mean of the four module scores. Differences between M0 and M1 tumors were assessed using the Mann– Whitney U test, with Benjamini–Hochberg correction applied to the four module-level comparisons.

### 2.9. Mediation Analysis

To investigate whether cg10057218 methylation statistically accounted for the association between metastatic status and GSDMB expression, a regression-based mediation analysis was performed. Metastatic status (M0 = 0, M1 = 1) was specified as the predictor (X), cg10057218 methylation β-value as the mediator (M), and GSDMB expression [log₂(RSEM + 1)] as the outcome (Y). Path coefficients were estimated using ordinary least-squares linear regression models with an intercept. The total association (c), direct association (c′), and indirect association (a × b) were estimated using unstandardized regression coefficients. Statistical inference for the indirect association was based primarily on 5,000 bootstrap resamples to obtain a percentile 95% confidence interval, with the Sobel test additionally reported as a complementary analysis. The proportion statistically accounted for by the indirect association was calculated as the ratio of the indirect association to the total association. Because metastatic status, DNA methylation, and gene expression were assessed using cross-sectional observational data, the mediation analysis was interpreted as a statistical decomposition of associations rather than evidence of temporal ordering or a causal mediation mechanism [22,23].

### 2.10. Canonical Molecular Score and Survival Analysis

To summarize the discriminatory information captured by the integrated multiomics model, a canonical molecular score was constructed using the 20 features with the largest absolute coefficients from the L2-regularized logistic regression model fitted to the full matched cohort. For each patient, the score was calculated as the weighted sum of standardized feature values according to Score = Σ(βᵢ × zᵢ), where βᵢ represents the L2-regularized logistic regression coefficient for feature i and zᵢ represents the corresponding standardized patient-level feature value. Higher scores indicated greater model-derived association with the M1 class.

Score distributions were compared between M0 and M1 tumors using the Mann– Whitney U test and across molecular subtypes using the Kruskal–Wallis test. The apparent discriminative performance of the full-cohort-derived score was summarized using ROC-AUC, sensitivity, and specificity. Because feature selection, coefficient estimation, and score evaluation were performed using the same cohort, these metrics were considered descriptive in-sample estimates and were not interpreted as measures of out-of-sample predictive performance. Generalizable predictive performance was instead assessed using the nested cross-validation framework described in Section 2.3. The score was additionally evaluated within the chromosomal instability (CIN) subgroup as an exploratory analysis of discrimination between M0 and M1 tumors within this molecular subtype.

The association between the canonical molecular score and overall survival was explored using Kaplan–Meier analysis. Patients with evaluable survival data were dichotomized into high- and low-score groups using the cohort median score as the cutoff, and survival distributions were compared using the log-rank test. Because this analysis was unadjusted and the molecular score was specifically derived to discriminate metastatic status, which is itself strongly associated with survival, the survival analysis was considered exploratory and was not interpreted as evidence of independent prognostic value.

### 2.11. Statistical Analysis

Comparisons between non-metastatic (M0) and metastatic (M1) tumors were performed using two-sided Mann–Whitney U tests for continuous variables. Categorical variables were compared using Fisher’s exact test for 2 × 2 contingency tables and Pearson’s chi-squared test for larger contingency tables. Unless otherwise specified, all statistical tests were two-sided, and statistical significance was defined as p < 0.05. Where multiple comparisons were performed, p-values were adjusted using the Benjamini– Hochberg false discovery rate procedure as described in the corresponding analyses.

## 3. Results

### 3.1. Cohort Characteristics

The final matched multiomics cohort comprised 518 patients with available RNA-seq and DNA methylation data and defined pathological metastasis status, including 436 M0 and 82 M1 cases, corresponding to an approximately 5.3:1 class ratio (Table 1). Patients with M1 disease were slightly younger than those with M0 disease, although the difference was not statistically significant (median age, 65 vs. 68 years; p = 0.074). As expected, metastatic disease was associated with more advanced locoregional tumor burden. T4 tumors were more frequent in the M1 group than in the M0 group (30.5% vs. 7.1%; p < 0.001), as was nodal involvement (N1/N2: 87.8% vs. 32.6%; p < 0.001). The chromosomal instability (CIN) molecular subtype was markedly enriched among M1 tumors (91.7% vs. 68.1%; p = 0.003), whereas the microsatellite instability (MSI) subtype was less frequent (3.3% vs. 15.1%). All-cause mortality was approximately threefold higher in patients with M1 disease than in those with M0 disease (43.9% vs. 14.9%; p < 0.001). Median follow-up for overall survival was also shorter in the M1 group (14 vs. 22 months; p < 0.001), consistent with the more aggressive clinical course of metastatic disease.

**Table 1.** Key clinical characteristics of the matched TCGA-COAD/READ multiomics cohort (n = 518) stratified by pathological metastasis status (M0 vs. M1). AJCC stage is omitted from this table because stage IV is definitional for M1 disease; the complete clinical breakdown is provided in Supplementary Table S1. Continuous variables: Mann– Whitney U test. Categorical variables: chi-squared or Fisher’s exact test.

| Variable | Total | M0 | M1 | p-value |
| --- | --- | --- | --- | --- |
| Patients, n | 518 | 436 | 82 |  |
| Age, median (min–max), years | 67.5 (31–90) | 68.0 (31–90) | 65.0 (35–87) | 0.074 |
| T4 primary tumor, n (%) | 56 (10.8%) | 31 (7.1%) | 25 (30.5%) | <0.001 |
| Nodal involvement (N1/N2), n (%) | 214 (41.3%) | 142 (32.6%) | 72 (87.8%) | <0.001 |
| CIN molecular subtype, n (%) | 281 (71.7%) | 226 (68.1%) | 55 (91.7%) | 0.003 |
| MSI molecular subtype, n (%) | 52 (13.3%) | 50 (15.1%) | 2 (3.3%) |  |
| Death (all-cause), n (%) | 101 (19.5%) | 65 (14.9%) | 36 (43.9%) | <0.001 |
| OS follow-up, median (min–max), months | 20.9 (0–148) | 22.0 (0–148) | 14.0 (0–82) | <0.001 |

### 3.2. Multiomics Integration Improves Metastasis Prediction

We compared five omics configurations across five classifiers (Figure 1). Somatic mutation and RPPA models showed the lowest overall discriminative performance. RNA-seq alone achieved moderate performance, with ROC-AUC values up to 0.731 ± 0.077, while DNA methylation reached a maximum ROC-AUC of 0.712 ± 0.047. The integration of RNA-seq and DNA methylation yielded the highest mean ROC-AUC in four of the five classifiers, with the best performance observed for linear SVM (0.787 ± 0.047), followed by Logistic Regression (0.775 ± 0.034), LightGBM (0.746 ± 0.048), and XGBoost (0.717 ± 0.044). Random Forest achieved a slightly higher mean ROC-AUC with RNA-seq alone than with the integrated data. Overall, these results suggest that the combined transcriptomic and DNA methylation representation provides complementary information for discriminating metastatic (M1) from non-metastatic (M0) tumors.

**Figure 1.**
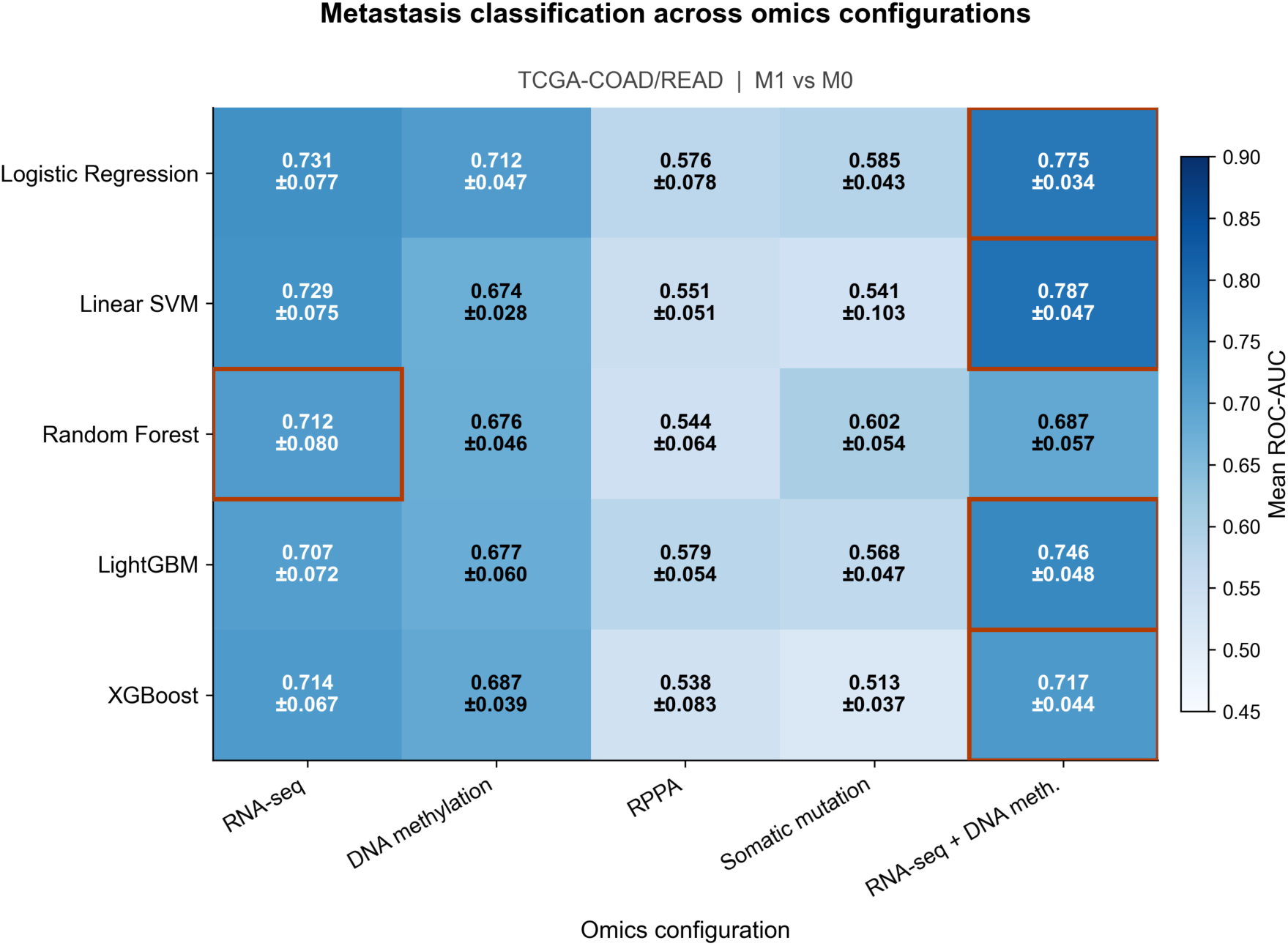
Cross-validated discriminative performance across omics configurations and machine-learning classifiers. The heatmap shows mean ROC-AUC ± standard deviation across the five outer folds of nested stratified cross-validation for RNA-seq, DNA methylation, RPPA, somatic mutation, and integrated RNA-seq + DNA methylation models. Outlined cells indicate the omics configuration with the highest mean ROC-AUC for each classifier. The integrated RNA-seq + DNA methylation configuration yielded the highest mean ROC-AUC in four of the five classifiers, with the highest overall performance observed for the linear SVM (0.787 ± 0.047).

### 3.3. Consensus Predictive Signature Across Four Models

To identify robust predictors, we extracted the top 20 features from each model based on absolute importance and identified those shared across all four models (Figure 2). Six features constituted the four-model consensus signature: ARC, C4orf23 (TRMT44), ASPDH, GPATCH3, cg12040555, and C13orf15. ARC ranked first in Logistic Regression (coefficient = +0.371) and XGBoost (mean |SHAP| = 0.286), and second in linear SVM (coefficient = +0.173). C4orf23, which encodes the putative tRNA methyltransferase TRMT44, was the highest-ranked feature in LightGBM. Directionality was consistent across all four classifiers for the six consensus features: ARC, ASPDH, and C13orf15 were associated with the M1 class, whereas C4orf23, GPATCH3, and cg12040555 were associated with the M0 class.

**Figure 2.**
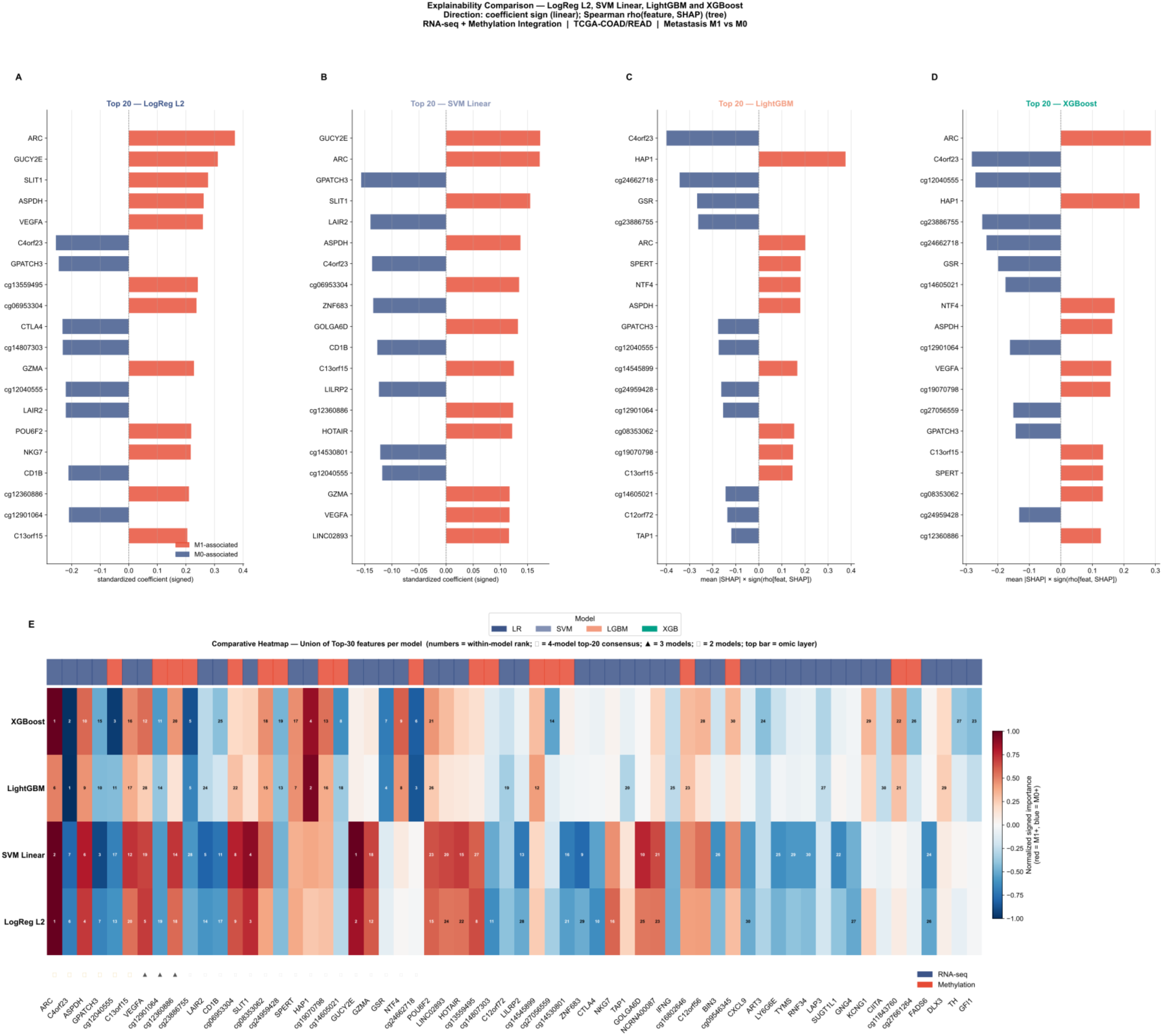
Consensus predictive signature across four classifiers. (A–D) Top 20 features ranked by absolute importance for Logistic Regression L2, linear SVM, LightGBM, and XGBoost. For the linear models, feature importance was based on the absolute standardized coefficient and direction on the coefficient sign. For the tree-based models, importance was quantified using mean absolute SHAP values, with direction determined by the sign of the Spearman correlation between feature values and their corresponding SHAP values across samples. Red indicates M1-associated features and blue indicates M0-associated features. (E) Comparative heatmap showing the union of the top 30 features from each model, with signed importance normalized within each model. Numbers indicate within-model rank. ★ indicates features present among the top 20 in all four models, ▴ in three models, and ◆ in two models. The top annotation indicates the omics layer (RNA-seq or DNA methylation).

Three additional features—VEGFA, cg12901064, and cg12360886, the latter annotated to GSDMB—appeared among the top 20 features in three of the four integrated models. Complementary cross-validation feature-importance stability analysis further highlighted GSDMB-associated methylation. Among methylation features, cg12360886 showed maximal stability, appearing among the top 50 features in all 15 fold–model combinations (stability = 1.000; mean rank = 7.73). A second GSDMB-associated probe, cg10057218, which was retained within the common 300-feature set used across the four integrated models, appeared among the top 50 features in 13 of 15 fold–model combinations (stability = 0.867; mean rank = 15.92). Thus, two independent CpG probes annotated to GSDMB were recurrently prioritized across complementary model-based analyses, motivating further investigation of the relationship between GSDMB methylation, gene expression, and metastatic status (Supplementary Table S2).

### 3.4. GSDMB Epigenetic Silencing in Metastatic Tumors

Given the recurrent prioritization of two GSDMB-associated CpG probes in the model-based and cross-validation feature-importance stability analyses, we next investigated their relationship with GSDMB expression. Among the 101 model-derived CpG probes, 86 had eligible single-gene CpG–gene pairs in the methylation–expression analysis. cg10057218 and cg12360886 exhibited the strongest and second-strongest inverse methylation–expression correlations, respectively, among these 86 model-derived pairs. cg10057218 methylation was inversely correlated with GSDMB expression (Spearman’s ρ = −0.589, raw p = 9.61 × 10⁻⁵⁰, BH-FDR = 1.05 × 10⁻⁴⁷), as was cg12360886 (ρ = −0.576, raw p = 3.46 × 10⁻⁴⁷, BH-FDR = 3.40 × 10⁻⁴⁵). Both associations therefore remained highly significant after correction across the full set of 19,150 CpG–gene correlation tests. Thus, two independent GSDMB-associated CpG probes were convergently prioritized by model-based analyses, cross-validation feature-importance stability, and methylation–expression association.

Both GSDMB-associated probes were significantly hypermethylated in M1 primary tumors compared with M0 primary tumors (Mann–Whitney U test, p < 0.0001 for both), with median β-values increasing from approximately 0.43 to 0.58 for cg10057218 and from 0.63 to 0.83 for cg12360886. Consistent with their inverse methylation– expression relationships, GSDMB expression was significantly lower in M1 tumors (Δlog₂ expression = −0.50, raw p = 4.18 × 10⁻⁴, BH-FDR = 0.036) (Figure 3). Together, the increased methylation of both GSDMB-associated CpGs in M1 tumors, their strong inverse correlations with GSDMB expression, and the concurrent reduction in GSDMB expression demonstrate a strong association between increased GSDMB-associated methylation and reduced GSDMB expression in M1 tumors. Somatic mutations in GSDMB were rare (6 patients, 1.2%), indicating that recurrent genetic alteration of GSDMB was uncommon in this cohort.

**Figure 3.**
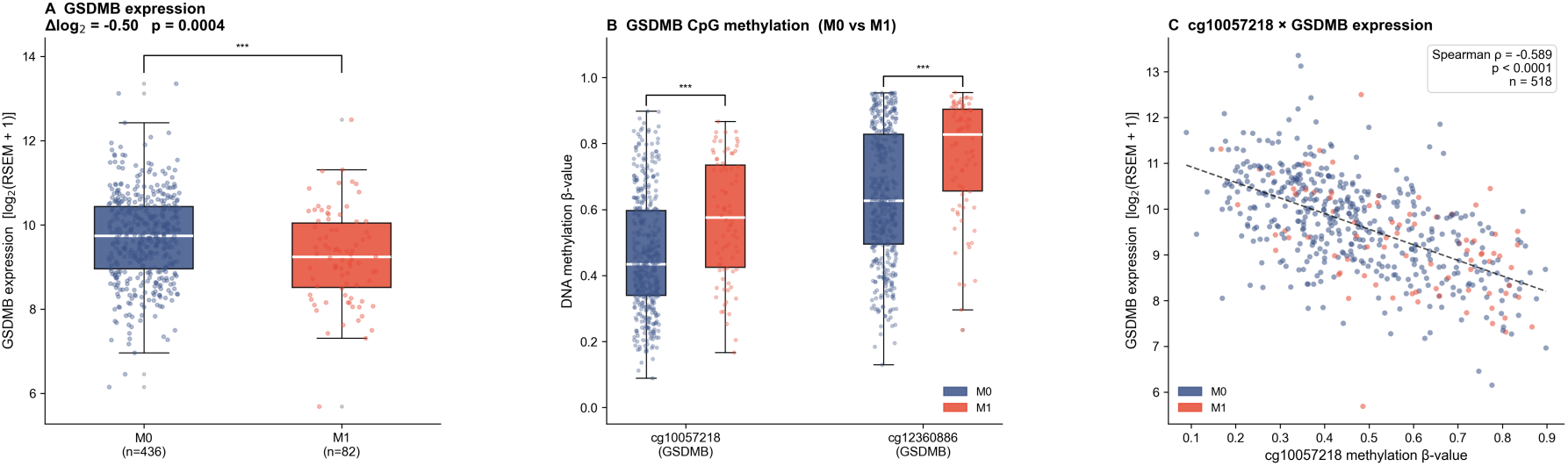
Multiomics profile of GSDMB in non-metastatic (M0) and metastatic (M1) colorectal tumors. (A) GSDMB expression [log₂(RSEM + 1)] in M0 vs. M1 tumors. (B) Methylation β-values of the GSDMB-associated CpG probes cg10057218 and cg12360886 in M0 vs. M1 tumors. (C) Scatter plot of cg10057218 methylation β-values and GSDMB expression (Spearman ρ = −0.589, p = 9.61 × 10⁻⁵⁰); the fitted regression line is shown for visualization. Group comparisons were performed using the Mann–Whitney U test; *** p < 0.001.

### 3.5. Functional Enrichment of M1-Associated Model-Derived Genes

Functional enrichment analysis of the M1-associated model-derived RNA-seq genes (ARC, ASPDH, GOLGA6D, GUCY2E, GZMA, HOTAIR, POU6F2, SLIT1, and VEGFA) identified Hedgehog signaling as the only significantly enriched MSigDB Hallmark pathway (FDR = 0.018), driven by SLIT1 and VEGFA. Given the small number of genes contributing to this enrichment, this finding should be interpreted as an over-representation signal rather than evidence of pathway activation.

Among genes with reduced expression in M1 tumors, Gene Ontology enrichment identified immune-related biological processes, including response to type II interferon (FDR = 2.08 × 10⁻⁹), T-cell activation (FDR = 1.23 × 10⁻⁶), and immunoglobulin-mediated immune response (FDR = 1.24 × 10⁻⁵). These results indicate enrichment of immune-related biological processes among genes with reduced expression in M1 tumors and motivated direct evaluation of immune-related transcriptional activity.

**Figure 4.**
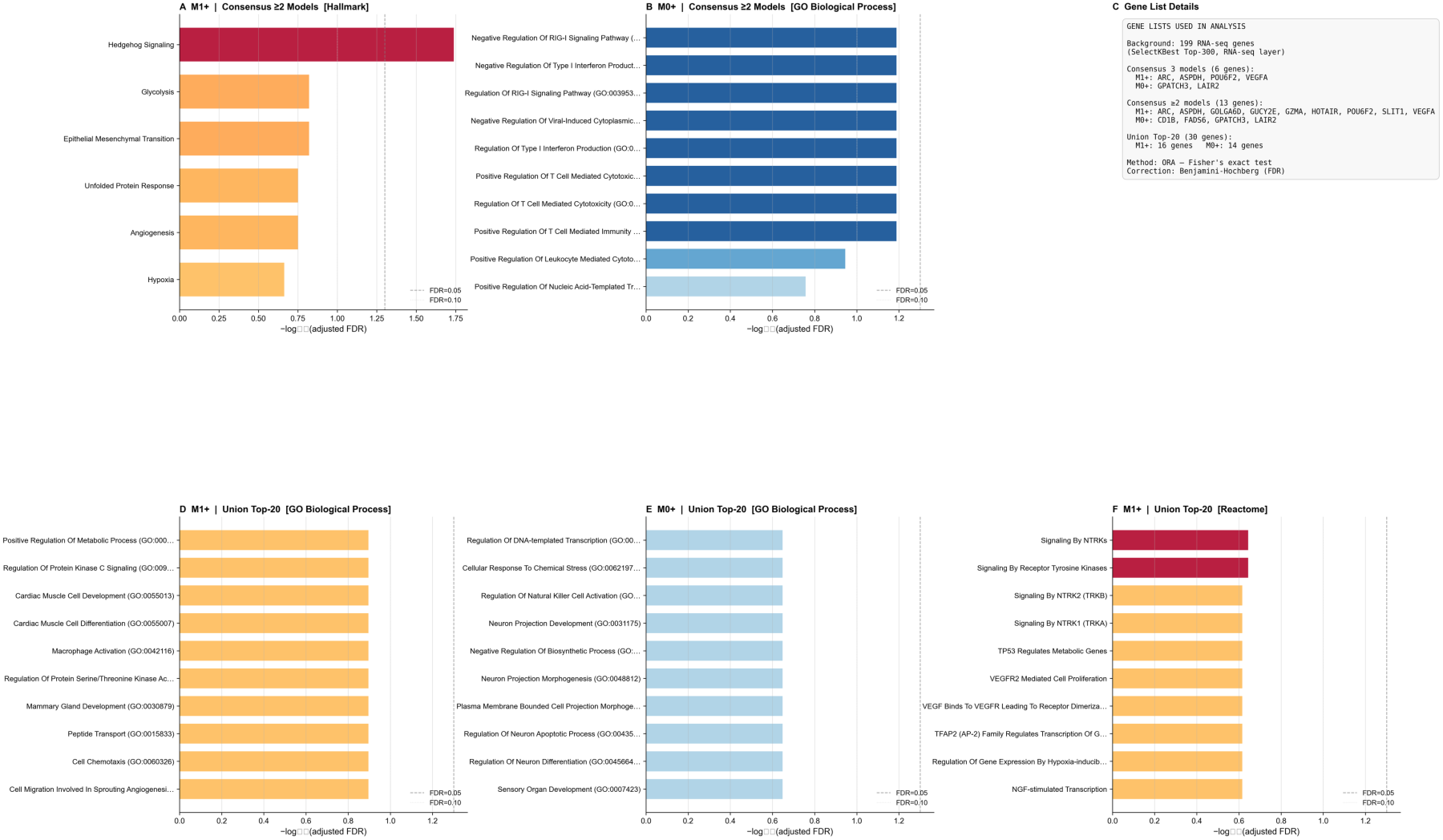
Functional enrichment of M1-associated model-derived RNA-seq genes. Over-representation analysis (ORA) against MSigDB Hallmark 2020 using nine M1-associated RNA-seq genes prioritized by the machine-learning models (ARC, ASPDH, GOLGA6D, GUCY2E, GZMA, HOTAIR, POU6F2, SLIT1, and VEGFA). Bubble size indicates the gene ratio, and color indicates −log₁₀(adjusted p-value). Hedgehog signaling was significantly enriched (FDR = 0.018), driven by SLIT1 and VEGFA.

### 3.6. Immune-Related Transcriptional Activity Is Reduced in M1 Tumors

To assess immune-related transcriptional differences between metastatic and non-metastatic tumors, we evaluated a 29-gene panel comprising cytotoxic T/NK, interferon-γ response, antigen-presentation, and immune checkpoint/Treg-associated modules. The composite immune-related transcriptional score was significantly lower in M1 than in M0 tumors (M1 = −0.340 ± 0.693 vs. M0 = +0.064 ± 0.750; Mann–Whitney p = 7.79 × 10⁻⁶; ROC-AUC = 0.656). All four functional modules were significantly reduced in M1 tumors, with the largest difference observed for the interferon-γ response module (Δ = −0.545; FDR = 5.38 × 10⁻⁷), followed by checkpoint/Treg-associated expression (Δ = −0.407; FDR = 1.78 × 10⁻⁴), cytotoxic T/NK activity (Δ = −0.350; FDR = 6.78 × 10⁻⁴), and antigen presentation (Δ = −0.312; FDR = 8.79 × 10⁻⁴). These findings indicate a coordinated reduction in immune-related transcriptional activity in M1 tumors (Figure 5).

**Figure 5.**
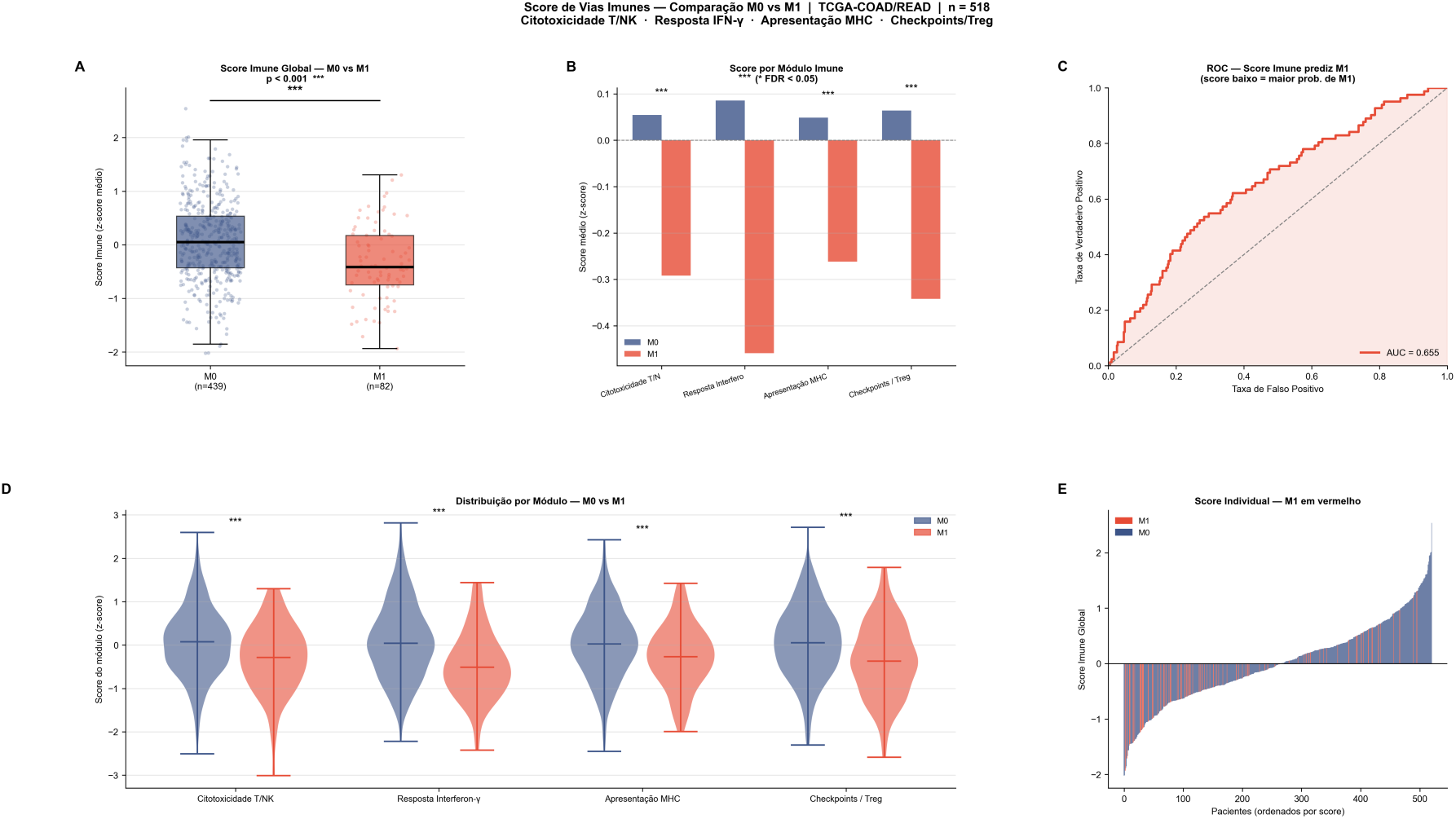
Immune-related transcriptional activity in non-metastatic (M0) and metastatic (M1) colorectal tumors. (A) Composite immune-related transcriptional score, calculated as the mean of four module scores, in M0 and M1 tumors; individual patient values are shown as jittered points. Values were calculated in the matched multiomics cohort (n = 518; M0 = 436, M1 = 82). (B) Mean module scores for M0 and M1 groups; asterisks indicate FDR-corrected significance (*** FDR < 0.001). (C) ROC curve for the composite immune score as a predictor of M1 status (AUC = 0.656; lower immune scores indicate greater association with M1 status). (D) Violin plots of module score distributions by metastatic status. (E) Individual patient scores ordered by composite immune score. (F) Per-gene differential expression (Δ = mean M1 z-score − mean M0 z-score), grouped by immune module. Significance was assessed using the Mann–Whitney U test with Benjamini–Hochberg correction for module-level comparisons.

### 3.7. Statistical Indirect Association Between Metastatic Status, GSDMB-Associated Methylation, and GSDMB Expression

Because cg10057218 exhibited the strongest inverse methylation–expression correlation with GSDMB among the eligible model-derived CpG–gene pairs (ρ = −0.589), while also showing high cross-validation feature-importance stability (13 of 15 fold–model combinations), this probe was prioritized for regression-based statistical mediation analysis.

To assess whether cg10057218 methylation statistically accounted for the association between metastatic status (M1 vs. M0) and GSDMB expression, we performed a regression-based mediation analysis with 5,000 bootstrap resamples for estimation of the indirect association. In this model, metastatic status (M0 = 0, M1 = 1) was specified as the predictor (X), cg10057218 methylation β-value as the mediator (M), and GSDMB log₂(RSEM + 1) expression as the outcome (Y). The total association between M1 status and GSDMB expression was significant (c = −0.451, p < 0.001). M1 status was associated with higher cg10057218 methylation (path a = +0.110, p = 3.28 × 10⁻⁷), and higher methylation was strongly associated with lower GSDMB expression after accounting for metastatic status (path b = −3.371, p = 4.14 × 10⁻⁴⁴). After accounting for cg10057218 methylation, the direct association between M1 status and GSDMB expression was substantially attenuated and no longer statistically significant (c′ = −0.080, p = 0.459). The estimated indirect association was −0.370, with a bootstrap 95% confidence interval excluding zero (95% CI, −0.519 to −0.224); the complementary Sobel test was also significant (p = 9.41 × 10⁻⁷). The indirect association corresponded to 82.1% of the total association. These findings indicate that cg10057218 methylation statistically accounted for a substantial proportion of the cross-sectional association between metastatic status and GSDMB expression (Figure 6).

**Figure 6.**
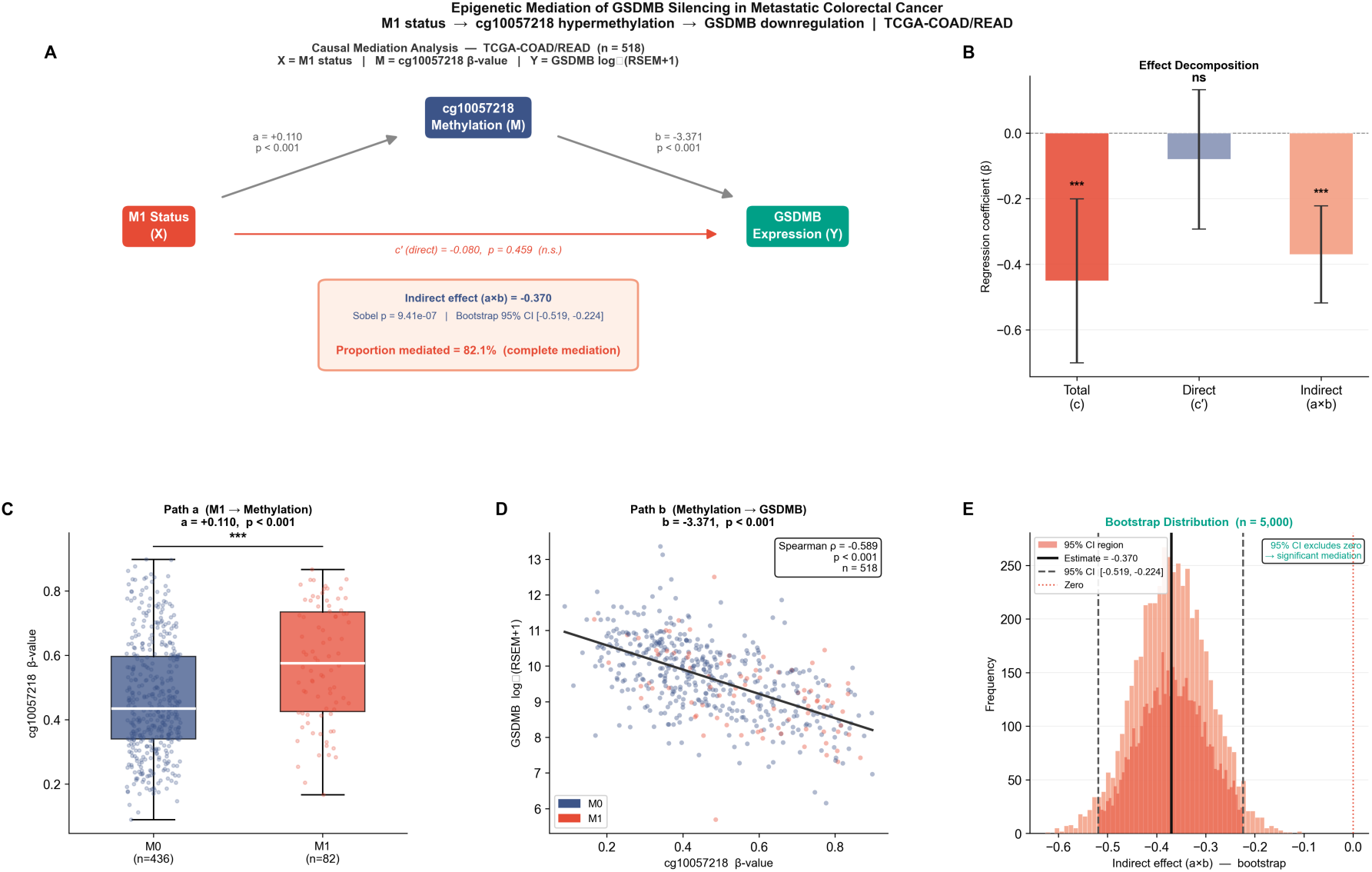
Regression-based statistical mediation analysis of the association between metastatic status, cg10057218 methylation, and GSDMB expression. Analysis was performed in the matched TCGA-COAD/READ multiomics cohort (n = 518; M0 = 436, M1 = 82), with metastatic status (M0 = 0, M1 = 1) as the predictor (X), cg10057218 β-value as the mediator (M), and GSDMB log₂(RSEM + 1) expression as the outcome (Y). (A) Path diagram showing unstandardized regression coefficients for paths a, b, and c′, together with the indirect effect (a × b). (B) Decomposition of the total (c), direct (c′), and indirect (a × b) effects. (C) cg10057218 β-values in M0 and M1 tumors. (D) Bivariate association between cg10057218 methylation and GSDMB expression; the fitted line and Spearman correlation are shown for visualization and are distinct from the multivariable regression coefficient for path b. (E) Bootstrap distribution of the indirect effect (5,000 resamples; 95% percentile CI [−0.519, −0.224]). The proportion mediated was 82.1%. Coefficients are unstandardized, and the mediation estimates represent statistical associations rather than evidence of causality.

Sensitivity analyses were performed to assess whether this indirect statistical association was influenced by molecular subtype composition or clinical covariates. When the analysis was restricted to CIN tumors (n = 281; M0 = 226, M1 = 55), M1 status remained associated with higher cg10057218 methylation (path a = +0.094, p = 6.63 × 10⁻⁴), while higher methylation remained strongly associated with lower GSDMB expression after accounting for M1 status (path b = −3.362, p = 5.43 × 10⁻²⁶). The indirect association remained different from zero (a × b = −0.316; bootstrap 95% CI, −0.533 to −0.115). A similar result was observed for cg12360886 (a × b = −0.273; bootstrap 95% CI, −0.451 to −0.098). In complete-case models adjusted for molecular subtype, age, sex, T stage, and N stage (n = 392), the indirect associations also remained different from zero for cg10057218 (a × b = −0.280; bootstrap 95% CI, −0.471 to −0.090) and cg12360886 (a × b = −0.249; bootstrap 95% CI, −0.409 to −0.094). Because the total association between M1 status and GSDMB expression was weak and non-significant in the CIN-restricted and covariate-adjusted analyses, the proportion-mediated statistic was not interpreted in these sensitivity analyses.

### 3.8. A Canonical Molecular Score Stratifies Tumors by Metastatic Status and Is Associated With Overall Survival

To summarize the discriminatory information captured by the integrated multiomics model, we constructed a canonical molecular score using the 20 features with the largest absolute coefficients from the L2-regularized logistic regression model. The score integrated 13 RNA-seq features (ARC, GUCY2E, SLIT1, ASPDH, VEGFA, C4orf23, GPATCH3, CTLA4, GZMA, LAIR2, NKG7, CD1B, and C13orf15) and seven DNA methylation features. M1 tumors had significantly higher canonical scores than M0 tumors (M1 = +1.659 ± 1.088; M0 = −0.312 ± 1.060; mean ± SD; Mann–Whitney U test, p = 2.97 × 10⁻³³; Cohen’s d = 1.849). The apparent in-sample ROC-AUC of the canonical score was 0.918 (sensitivity = 0.902; specificity = 0.821 at the Youden-optimal threshold). Because the score was derived and evaluated in the same cohort, this value represents descriptive in-sample discrimination rather than out-of-sample predictive performance. By comparison, the corresponding logistic regression model achieved a nested cross-validated ROC-AUC of 0.781 ± 0.046.

Canonical score distributions differed significantly across molecular subtypes (Kruskal–Wallis H = 27.97, p = 3.68 × 10⁻⁶), with higher scores observed in CIN tumors than in MSI and GS tumors. Because CIN was more frequent among M1 tumors, we additionally evaluated the score within the CIN subgroup. Among CIN tumors (n = 281; M0 = 226, M1 = 55), M1 tumors retained significantly higher scores than M0 tumors (Mann–Whitney U test, p = 4.28 × 10⁻¹⁸), with an apparent in-sample ROC-AUC of 0.877. This exploratory within-subtype analysis indicates that the association between the canonical score and metastatic status was retained within CIN tumors, although the estimate remains in-sample and should not be interpreted as externally validated discrimination.

In unadjusted Kaplan–Meier analysis, patients with high molecular scores, defined using the median score among patients with evaluable survival data, had significantly shorter overall survival than patients with low scores (log-rank χ² = 27.38, p = 1.67 × 10⁻⁷). However, because the canonical score was derived to discriminate metastatic status and the survival analysis was not adjusted for M-stage or other clinicopathological covariates, this association should not be interpreted as evidence of independent prognostic value. Rather, the survival analysis provides exploratory evidence that the molecular pattern captured by the score is also associated with overall survival (Figure 7).

**Figure 7.**
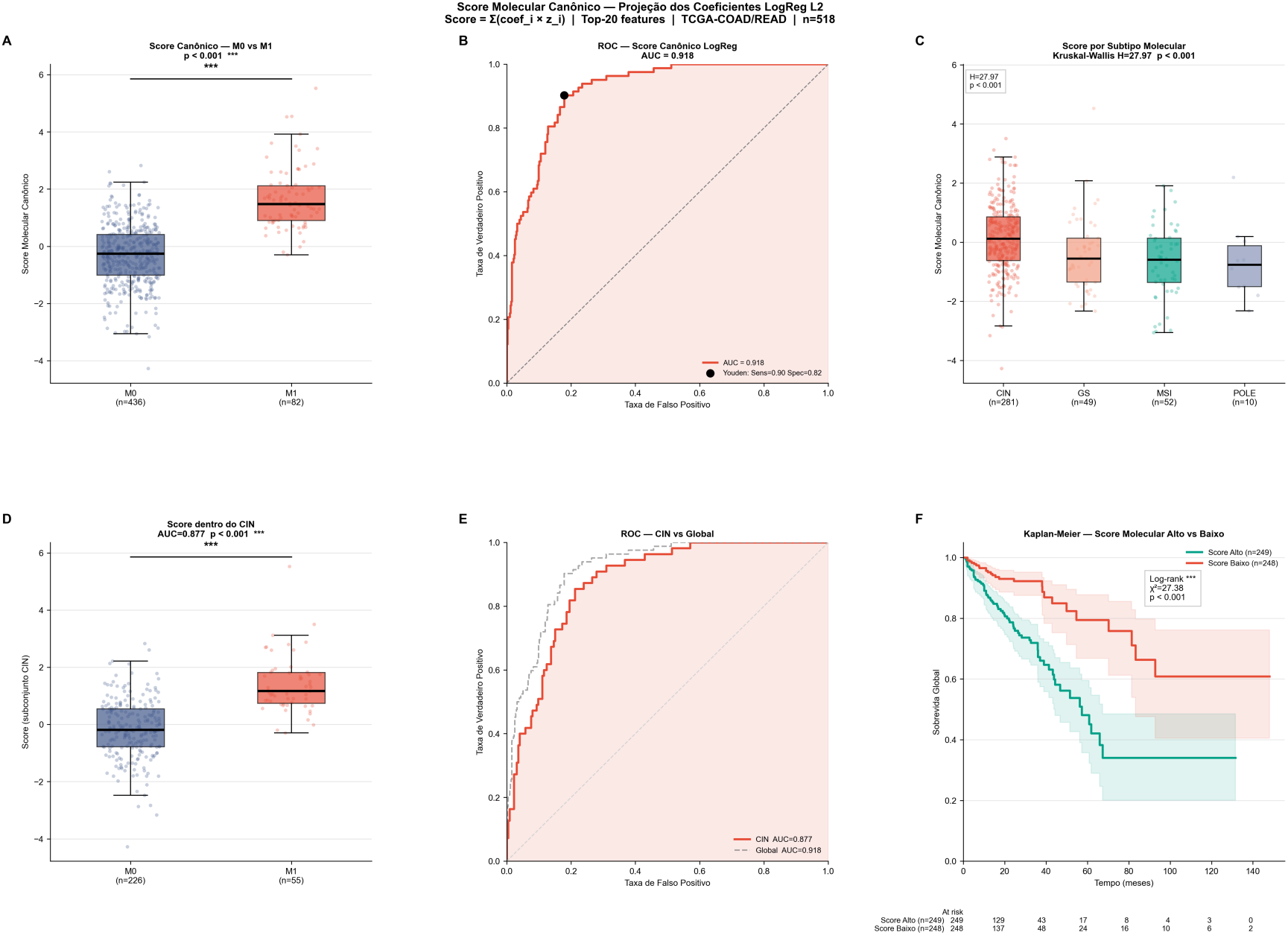
Canonical molecular score derived from the L2-regularized logistic regression model. (A) Distribution of the canonical molecular score in M0 and M1 tumors (Mann–Whitney U test). (B) In-sample ROC curve for discrimination between M0 and M1 tumors using the full-cohort-derived canonical score (ROC-AUC = 0.918); because the score was derived and evaluated in the same cohort, this represents apparent discrimination, whereas the corresponding logistic regression model achieved a nested cross-validated ROC-AUC of 0.781 ± 0.046. (C) Canonical molecular score distributions across molecular subtypes (Kruskal–Wallis test). (D) Canonical molecular score in M0 and M1 tumors within the CIN subtype (Mann–Whitney U test). (E) In-sample ROC curve within the CIN subgroup (ROC-AUC = 0.877). (F) Unadjusted Kaplan–Meier analysis of overall survival for high-versus low-score groups defined by the median canonical score among patients with evaluable survival data; survival distributions were compared using the log-rank test. The survival analysis was exploratory and was not adjusted for metastatic status or other clinicopathological covariates.

## 4. Discussion

This study demonstrates that the integration of RNA-seq transcriptomics and DNA methylation data improves the discrimination of CRC metastatic status compared with individual omics layers across most evaluated classifiers. Integrated RNA-seq and DNA methylation models achieved the highest ROC-AUC in four of the five classifiers, with the best overall performance observed for linear SVM (ROC-AUC = 0.787 ± 0.047). Random Forest represented the exception, showing slightly better performance with RNA-seq alone than with the integrated data. Overall, these findings suggest that transcriptomic and epigenomic features provide complementary molecular information for distinguishing M1 from M0 tumors. This observation is consistent with the broader multiomics literature, in which integration of molecular layers can improve cancer classification by capturing complementary biological signals [5,6].

Beyond overall model performance, explainability analyses identified a set of molecular features that were consistently prioritized across classifiers with distinct learning strategies. Six features—ARC, ASPDH, C13orf15, C4orf23, GPATCH3, and cg12040555—ranked among the top 20 predictors across Logistic Regression, SVM, LightGBM, and XGBoost, with consistent directions of association. Their convergence across linear and tree-based models supports the robustness of the predictive signal, although their biological roles in metastatic CRC cannot be inferred from model importance alone. Rather than establishing individual drivers of metastasis, these features provide a compact set of candidates for subsequent biological investigation.

Among the methylation-associated findings, GSDMB emerged as a particularly notable candidate. Two CpG probes annotated to GSDMB, cg10057218 and cg12360886, showed strong inverse correlations with GSDMB expression, with cg10057218 exhibiting the strongest association (ρ = −0.589). cg10057218 methylation was also significantly higher in M1 tumors, whereas GSDMB expression was reduced. Regression-based statistical mediation further showed a significant indirect association between metastatic status and GSDMB expression through cg10057218 methylation, with an estimated 82.1% proportion mediated. Together, these findings support a statistical relationship among metastatic status, increased cg10057218 methylation, and reduced GSDMB expression. Given the involvement of GSDMB in gasdermin-mediated cell death and tumor biology [15–17], its epigenetic regulation may represent a biologically relevant feature of metastatic CRC. However, because these analyses are based on cross-sectional observational data, they do not establish that cg10057218 methylation causally drives GSDMB repression or metastatic progression. Functional studies will therefore be required to determine the mechanistic consequences of this methylation–expression relationship.

The reduced immune-related transcriptional activity observed in M1 tumors provides an additional biological context for the molecular differences associated with metastatic disease. All four evaluated immune modules—cytotoxic T/NK activity, interferon-γ response, antigen presentation, and checkpoint/Treg-associated expression— were significantly reduced in M1 tumors, with the largest difference observed for the interferon-γ response module. These findings indicate a broad reduction in immune-associated transcriptional programs in metastatic tumors and are consistent with alterations in tumor–immune interactions accompanying CRC progression. In this context, the concurrent reduction in GSDMB expression is noteworthy given the involvement of gasdermin family proteins in inflammatory cell death and their potential influence on antitumor immune responses [15–17]. However, the present analyses do not establish a mechanistic relationship between GSDMB repression and reduced immune-related transcriptional activity. Rather, these findings identify concurrent epigenetic, transcriptional, and immune-associated features of M1 disease whose potential biological relationships warrant investigation in experimental models.

Functional enrichment analyses provided additional biological context for the molecular differences associated with metastatic status. Among M1-associated model-derived RNA-seq features, Hedgehog signaling was significantly enriched (FDR = 0.018), with SLIT1 and VEGFA contributing to this association. Given the established roles of Hedgehog-related signaling and VEGFA in tumor progression, angiogenesis, and metastatic processes, these findings identify potentially relevant biological programs associated with M1 disease. Complementarily, genes showing lower expression in M1 tumors were strongly enriched for immune-related processes, including interferon-γ response, T-cell activation, and B-cell-mediated immunity. These enrichment results are consistent with the independently observed reduction in immune-related transcriptional scores in M1 tumors, providing convergent evidence of altered immune-associated transcriptional programs in metastatic disease. Because these analyses are based on gene-set over-representation, however, pathway enrichment should not be interpreted as direct evidence of pathway activation or suppression.

The canonical molecular score provided a complementary summary of the discriminatory information captured by the integrated model. The 20-feature score showed strong separation between M0 and M1 tumors in the derivation cohort (in-sample ROC-AUC = 0.918), while the corresponding Logistic Regression model achieved a more modest ROC-AUC of 0.781 ± 0.046 under nested cross-validation. This difference underscores the importance of distinguishing apparent discrimination from cross-validated model performance when interpreting molecular scores derived and evaluated within the same cohort. The score also retained substantial in-sample discrimination within the CIN subgroup (ROC-AUC = 0.877), suggesting that its association with metastatic status was not solely attributable to differences across major molecular subtypes. Higher scores were additionally associated with shorter overall survival in unadjusted Kaplan–Meier analysis (log-rank p = 1.67 × 10⁻⁷). However, because the survival analysis was not adjusted for metastatic status or other clinicopathological covariates, this association should not be interpreted as evidence of independent prognostic value. Rather, the canonical score provides a compact representation of the molecular features associated with metastatic status and illustrates their potential relationship with clinically relevant outcomes.

Several limitations should be acknowledged. First, the analysis is retrospective and cross-sectional; therefore, causal relationships between the identified molecular features and metastatic progression cannot be established. Second, the imbalance between M0 and M1 cases (436 vs. 82) may affect the precision and stability of metastatic-class discrimination despite the use of class-balancing strategies and stratified nested cross-validation. Third, the analysis was restricted to primary tumors, and their molecular profiles may not fully represent those of established metastatic lesions or the dynamic changes occurring during metastatic progression. Fourth, the absence of external validation limits the generalizability of the identified multiomics signature, and replication in independent cohorts with comparable transcriptomic, methylation, and metastatic-status information will be necessary. Fifth, although RNA-seq and DNA methylation integration improved discrimination across most classifiers, the magnitude of improvement varied by model, emphasizing the complexity and heterogeneity of the molecular features associated with metastatic disease. Finally, the canonical molecular score was derived and evaluated within the same cohort, making its ROC-AUC an apparent in-sample estimate, while its association with overall survival was assessed without adjustment for metastatic status or other clinicopathological covariates. Independent validation and multivariable survival analyses will therefore be required to establish its potential clinical utility and prognostic relevance.

### Conclusions

This study presents an interpretable multiomics machine learning framework integrating transcriptomic and DNA methylation data to characterize molecular features associated with metastatic status in colorectal cancer. RNA-seq and DNA methylation integration improved discrimination between M0 and M1 tumors across most evaluated classifiers and identified a robust set of molecular features consistently prioritized across distinct modeling approaches. Subsequent analyses revealed coordinated transcriptional, epigenetic, and immune-related differences associated with M1 disease. In particular, increased cg10057218 methylation was strongly associated with reduced GSDMB expression and showed a significant statistical mediation of the relationship between metastatic status and GSDMB expression, identifying this methylation–expression relationship as a candidate feature for further mechanistic investigation. M1 tumors also exhibited reduced immune-related transcriptional activity and enrichment of distinct biological processes associated with metastatic disease. Together, these findings demonstrate the value of interpretable multiomics integration for identifying complementary molecular features associated with CRC metastasis and provide candidates for validation in independent cohorts and experimental models.

## Author contributions

**Matheus da Silveira Costa:** Conceptualization, Methodology, Data Curation, Formal Analysis, Software, Visualization, Investigation, Writing – Original Draft. **Henrique Izaias Marcelo:** Review & Editing. **Gabriel Albanese Kafouri:** Review & Editing. **Vinicius de Camargo:** Conceptualization, Supervision, Writing – Review & Editing.

## Data and Code Availability

TCGA-COAD/READ data are publicly available through cBioPortal. All analysis code and supporting computational outputs used in this study are publicly available in the GSDMB-CRC-Multiomics GitHub repository: GSDMB-CRC-Multiomics.

