## Supplementary Table S1 for "Interpretable Multiomics Machine Learning Identifies GSDMB-Associated Epigenetic Repression and Reduced Immune Activity in Metastatic Colorectal Cancer"

| Variable | Total | M0 | M1 | p-value |
|---|---|---|---|---|
| **N** | 518 | 436 | 82 | |
| **Age at diagnosis, median (min–max)** | 67.5 (31–90) | 68.0 (31–90) | 65.0 (35–87) | 74 |
| **Sex — Male, n (%)** | 271 (52.3%) | 224 (51.4%) | 47 (57.3%) | 337 |
| **Sex — Female, n (%)** | 247 (47.7%) | 212 (48.6%) | 35 (42.7%) | |
| **AJCC stage — I, n (%)** | 92 (17.9%) | 92 (21.2%) | 0 (0.0%) | <0.001 |
| **AJCC stage — II, n (%)** | 200 (38.8%) | 200 (46.1%) | 0 (0.0%) | |
| **AJCC stage — III, n (%)** | 141 (27.4%) | 141 (32.5%) | 0 (0.0%) | |
| **AJCC stage — IV, n (%)** | 82 (15.9%) | 1 (0.2%) | 81 (100.0%) | |
| **T stage — T1, n (%)** | 17 (3.3%) | 17 (3.9%) | 0 (0.0%) | <0.001 |
| **T stage — T2, n (%)** | 89 (17.2%) | 87 (20.0%) | 2 (2.4%) | |
| **T stage — T3, n (%)** | 356 (68.7%) | 301 (69.0%) | 55 (67.1%) | |
| **T stage — T4, n (%)** | 56 (10.8%) | 31 (7.1%) | 25 (30.5%) | |
| **N stage — N0, n (%)** | 304 (58.7%) | 294 (67.4%) | 10 (12.2%) | <0.001 |
| **N stage — N1, n (%)** | 122 (23.6%) | 88 (20.2%) | 34 (41.5%) | |
| **N stage — N2, n (%)** | 92 (17.8%) | 54 (12.4%) | 38 (46.3%) | |
| **Molecular subtype — CIN, n (%)** | 281 (71.7%) | 226 (68.1%) | 55 (91.7%) | 3 |
| **Molecular subtype — MSI, n (%)** | 52 (13.3%) | 50 (15.1%) | 2 (3.3%) | |
| **Molecular subtype — GS, n (%)** | 49 (12.5%) | 46 (13.9%) | 3 (5.0%) | |
| **Molecular subtype — POLE, n (%)** | 10 (2.6%) | 10 (3.0%) | 0 (0.0%) | |
| **New tumor event after initial treatment — Yes, n (%)** | 86 (20.5%) | 61 (17.1%) | 25 (39.7%) | <0.001 |
| **Death (OS_STATUS = Deceased), n (%)** | 101 (19.5%) | 65 (14.9%) | 36 (43.9%) | <0.001 |
| **OS follow-up, months — median (min–max)** | 20.9 (0–148) | 22.0 (0–148) | 14.0 (0–82) | <0.001 |
| **Race — White, n (%)** | 235 (80.5%) | 198 (81.1%) | 37 (77.1%) | 103 |
| **Race — Black or African American, n (%)** | 45 (15.4%) | 34 (13.9%) | 11 (22.9%) | |
| **Race — Asian, n (%)** | 12 (4.1%) | 12 (4.9%) | 0 (0.0%) | |
