## Supplementary Table S2 for "Interpretable Multiomics Machine Learning Identifies GSDMB-Associated Epigenetic Repression and Reduced Immune Activity in Metastatic Colorectal Cancer"

stability_methylation_metastasis

| Feature | Frequenc | Mean_Rank | Mean_Importance | Stabilit | Layer | Combined_Score | Norm_Importance | Rank_Within_Laye | Outcome |
|---|---|---|---|---|---|---|---|---|---|
| cg12360886 | 5 | 3.0 | 0.025846098100486582 | 1.0 | Metilaçk | 0.824892672025713 | 0.9163089906752371 | 1 | metastasis |
| cg10057218 | 5 | 10.6 | 0.018104309375643026 | 1.0 | Metilaçk | 0.7004134352130057 | 0.6105735196755363 | 2 | metastasis |
| cg14605021 | 5 | 15.0 | 0.018105166406393876 | 1.0 | Metilaçk | 0.6956822095494857 | 0.6106073651649524 | 3 | metastasis |
| cg09721659 | 5 | 12.8 | 0.017805137724311447 | 1.0 | Metilaçk | 0.6917508879453791 | 0.5908804477406358 | 4 | metastasis |
| cg24030449 | 5 | 14.0 | 0.017559236927872396 | 1.0 | Metilaçk | 0.6900476593084404 | 0.5890477532503567 | 5 | metastasis |
| cg15901783 | 5 | 14.0 | 0.016382083465927 | 1.0 | Metilaçk | 0.6701502374039058 | 0.542723013568574 | 6 | metastasis |
| cg14458615 | 5 | 26.2 | 0.014281089009044042 | 1.0 | Metilaçk | 0.6452292683432557 | 0.45958775722261686 | 7 | metastasis |
| cg04576855 | 5 | 29.4 | 0.012199016061171717 | 1.0 | Metilaçk | 0.619788212959779 | 0.37736421863785996 | 8 | metastasis |
| cg24471894 | 4 | 10.25 | 0.019519776514304785 | 0.8 | Metilaçk | 0.6177195393226721 | 0.6684725384829809 | 9 | metastasis |
| cg09508556 | 5 | 28.6 | 0.011592339412759908 | 1.0 | Metilaçk | 0.6127784384501831 | 0.3534058058447545 | 10 | metastasis |
| cg17018527 | 5 | 42.4 | 0.010960127044010923 | 1.0 | Metilaçk | 0.6031398592244619 | 0.3284385469782096 | 11 | metastasis |
| cg22156632 | 5 | 46.4 | 0.010629522574872615 | 1.0 | Metilaçk | 0.5988341465704747 | 0.3153824578259146 | 12 | metastasis |
| cg09546345 | 5 | 44.2 | 0.010495102318457118 | 1.0 | Metilaçk | 0.5974469759814119 | 0.3100739907344899 | 13 | metastasis |
| cg03271907 | 5 | 54.8 | 0.008960806141440307 | 1.0 | Metilaçk | 0.5784911590957686 | 0.24968976568362522 | 14 | metastasis |
| cg00932276 | 5 | 65.8 | 0.008457490214506402 | 1.0 | Metilaçk | 0.5718758592231846 | 0.22960549082378912 | 15 | metastasis |
| cg16802646 | 4 | 23.75 | 0.014946298847232826 | 0.8 | Metilaçk | 0.5538384263536542 | 0.48585872757615334 | 16 | metastasis |
| cg25957124 | 4 | 30.75 | 0.012953384373555117 | 0.8 | Metilaçk | 0.5284458347690806 | 0.4071554072355181 | 17 | metastasis |
| cg25151295 | 4 | 35.0 | 0.011677469950189166 | 0.8 | Metilaçk | 0.5125857832658823 | 0.3587674257010892 | 18 | metastasis |
| cg07393322 | 1 | 1.0 | 0.02796531126234445 | 0.2 | Metilaçk | 0.4999999881525291 | 0.9999999605084305 | 19 | metastasis |
| cg24662718 | 4 | 38.0 | 0.010338760693763702 | 0.8 | Metilaçk | 0.4962981495080966 | 0.3038998146053485 | 20 | metastasis |
| cg08700306 | 4 | 55.75 | 0.009733578487468884 | 0.8 | Metilaçk | 0.4875342949179672 | 0.2800002194789245 | 21 | metastasis |
| cg16522484 | 4 | 56.25 | 0.009464531811981276 | 0.8 | Metilaçk | 0.4843059929838266 | 0.26937514400722395 | 22 | metastasis |
| cg00904483 | 4 | 53.0 | 0.008845004148320061 | 0.8 | Metilaçk | 0.47717641097959035 | 0.2449090242529554 | 23 | metastasis |
| cg17105163 | 4 | 66.5 | 0.007792711545979374 | 0.8 | Metilaçk | 0.46396886432344774 | 0.20335233786828258 | 24 | metastasis |
| cg18675299 | 4 | 64.5 | 0.007852486663894692 | 0.8 | Metilaçk | 0.462397826274244 | 0.19781463719913517 | 25 | metastasis |
| cg19954000 | 3 | 17.333333333333332 | 0.014286084254743 75 | 0.6 | Metilaçk | 0.4488448360530302 | 0.4597858171464643 | 26 | metastasis |
| cg27486427 | 3 | 21.666666666666668 | 0.014180398151509116 | 0.6 | Metilaçk | 0.4455071615300387 | 0.4556121070609135 | 27 | metastasis |
| cg24441911 | 3 | 20.333333333333332 | 0.013386346351715503 | 0.6 | Metilaçk | 0.43665112657928 | 0.42425375526426673 | 28 | metastasis |
| cg17740645 | 3 | 26.333333333333332 | 0.012742188973094268 | 0.6 | Metilaçk | 0.4289615639940847 | 0.3988149694111 7667 | 29 | metastasis |
| cg23886755 | 3 | 27.666666666666668 | 0.012542813271022446 | 0.6 | Metilaçk | 0.4242591371940315 | 0.3909413100266168 | 30 | metastasis |
| cg17065601 | 3 | 29.333333333333332 | 0.012310304055671378 | 0.6 | Metilaçk | 0.42112115345380924 | 0.38175915620134226 | 31 | metastasis |
| cg21305265 | 3 | 29.0 | 0.012280350950387 63 | 0.6 | Metilaçk | 0.4208395449658228 | 0.38057626106385384 | 32 | metastasis |
| cg21968169 | 2 | 9.0 | 0.019254089995456327 | 0.4 | Metilaçk | 0.41679404826178525 | 0.6559801608725506 | 33 | metastasis |
| cg12204727 | 3 | 36.0 | 0.011832359009178166 | 0.6 | Metilaçk | 0.41427071226707207 | 0.36288435620555565 | 34 | metastasis |
| cg12040555 | 3 | 33.333333333333336 | 0.011004758039029167 | 0.6 | Metilaçk | 0.40488557122381 82 | 0.3302010950179 0534 | 35 | metastasis |
| cg01324261 | 3 | 35.333333333333336 | 0.010882417540490633 | 0.6 | Metilaçk | 0.403115490173079 | 0.325369676723719 | 36 | metastasis |
| cg26256793 | 3 | 43.0 | 0.010713019837502996 | 0.6 | Metilaçk | 0.4001494232169879 | 0.3186798955717779 | 37 | metastasis |
| cg04201526 | 3 | 46.666666666666664 | 0.010399629908178621 | 0.6 | Metilaçk | 0.3960889481987375 | 0.3063030354135 8526 | 38 | metastasis |
| cg27056559 | 3 | 49.333333333333336 | 0.009536832473908128 | 0.6 | Metilaçk | 0.3856409362525555 | 0.2722225123049357 | 39 | metastasis |
| cg25227148 | 3 | 55.666666666666664 | 0.008502914064582074 | 0.6 | Metilaçk | 0.3736800500085586 | 0.2337688441204899 | 40 | metastasis |
| cg00873037 | 3 | 55.0 | 0.008303791092 69819 | 0.6 | Metilaçk | 0.3713429782057 4114 | 0.22590516544770853 | 41 | metastasis |
| cg02947222 | 3 | 53.333333333333336 | 0.008303038150697549 | 0.6 | Metilaçk | 0.3707327625209 5283 | 0.22350593641953612 | 42 | metastasis |
| cg17353431 | 1 | 4.0 | 0.02126031273772852 | 0.2 | Metilaçk | 0.3605827136879842 | 0.735209045626614 | 43 | metastasis |
| cg02887841 | 1 | 6.0 | 0.02200909174924171 | 0.2 | Metilaçk | 0.3580052797596661 | 0.7647735039607919 | 44 | metastasis |
| cg04317399 | 2 | 15.5 | 0.014737023892045717 | 0.4 | Metilaçk | 0.3553994514661253 | 0.4775941311497 1044 | 45 | metastasis |
| cg05955301 | 3 | 73.33333333333333 | 0.007060558679270625 | 0.6 | Metilaçk | 0.3550221388 0049545 | 0.1744385194761 8064 | 46 | metastasis |
| cg24959428 | 2 | 15.5 | 0.014602357810977 56 | 0.4 | Metilaçk | 0.3538039989982981 | 0.4722759562589 5323 | 47 | metastasis |
| cg04037952 | 3 | 80.0 | 0.006743801644378889 | 0.6 | Metilaçk | 0.3510479100 6540675 | 0.16192924754312543 | 48 | metastasis |
| cg11175477 | 3 | 72.66666666666667 | 0.006713461849161023 | 0.6 | Metilaçk | 0.35093425418117763 | 0.1607310735149 3604 | 49 | metastasis |
| cg08541219 | 2 | 18.0 | 0.014268322460153203 | 0.4 | Metilaçk | 0.3482279336483676 | 0.459005392862 9799 | 50 | metastasis |
| cg17066382 | 3 | 75.0 | 0.006425873291942735 | 0.6 | Metilaçk | 0.3474437039579214 | 0.1493737500351767 | 51 | metastasis |
| cg04297093 | 2 | 28.0 | 0.014432369440075423 | 0.4 | Metilaçk | 0.3465654063347873 | 0.4655828487021645 | 52 | metastasis |
| cg09829319 | 3 | 81.33333333333333 | 0.006242728870709017 | 0.6 | Metilaçk | 0.345071476619089 | 0.14214108940508048 | 53 | metastasis |
| cg13340269 | 2 | 24.5 | 0.013924195510804815 | 0.4 | Metilaçk | 0.3416099026155701 | 0.4458892178688938 | 54 | metastasis |
| cg23539753 | 1 | 7.0 | 0.020900650860572093 | 0.2 | Metilaçk | 0.341301630088 1567 | 0.7210054336271889 | 55 | metastasis |
| cg05874450 | 2 | 43.0 | 0.014046812917606184 | 0.4 | Metilaçk | 0.3396464395011678 | 0.4503366165190442 | 56 | metastasis |
| cg11843769 | 1 | 5.0 | 0.019859008474013527 | 0.2 | Metilaçk | 0.3372941358269461 | 0.6798693409787092 | 57 | metastasis |
| cg18401406 | 1 | 3.0 | 0.018407456880351215 | 0.2 | Metilaçk | 0.336783587123188 | 0.6225452904106267 | 58 | metastasis |
| cg17414107 | 2 | 28.5 | 0.013587246802111735 | 0.4 | Metilaçk | 0.3364359498229 0026 | 0.4321876293531704 | 59 | metastasis |
| cg22220722 | 2 | 23.0 | 0.013113190156967703 | 0.4 | Metilaçk | 0.332373249862125 | 0.4134603884293 0555 | 60 | metastasis |
| cg14424530 | 2 | 24.0 | 0.013058285 5250 7708 | 0.4 | Metilaçk | 0.33138943550379846 | 0.4112981183459 8483 | 61 | metastasis |
| cg07149772 | 2 | 26.0 | 0.012412431888793696 | 0.4 | Metilaçk | 0.3231451107920 9907 | 0.38579234461563877 | 62 | metastasis |
| cg03853987 | 1 | 6.0 | 0.018608437259 50501 | 0.2 | Metilaçk | 0.3177161248749117 | 0.6304823210116104 | 63 | metastasis |
| cg03361068 | 1 | 5.0 | 0.018094958332 63062 | 0.2 | Metilaçk | 0.31636460302 683425 | 0.610204232311 0032 | 64 | metastasis |
| cg16616769 | 2 | 29.0 | 0.011749721322804059 | 0.4 | Metilaçk | 0.31455292594 92004 | 0.35962086427511264 | 65 | metastasis |
| cg17274742 | 1 | 7.0 | 0.018417928327083413 | 0.2 | Metilaçk | 0.3118876472829408 | 0.6229588242764894 | 66 | metastasis |
| cg19139277 | 2 | 33.0 | 0.011517728586748637 | 0.4 | Metilaçk | 0.31102008481254645 | 0.3504591062379 | 67 | metastasis |
| cg08081407 | 2 | 34.0 | 0.011478583123940265 | 0.4 | Metilaçk | 0.310388006144294 | 0.34891240143336094 | 68 | metastasis |
| cg08353062 | 2 | 35.5 | 0.011350648813716436 | 0.4 | Metilaçk | 0.30863771142513385 | 0.34380086456779774 | 69 | metastasis |
| cg13565157 | 1 | 7.0 | 0.017283548078135057 | 0.2 | Metilaçk | 0.2984481103684967 | 0.578160367894989 | 70 | metastasis |
| cg17496788 | 2 | 34.5 | 0.010460998573445133 | 0.4 | Metilaçk | 0.2982519569128808 | 0.30872718031993135 | 71 | metastasis |
| cg14289511 | 2 | 39.5 | 0.009996897646643917 | 0.4 | Metilaçk | 0.29205800350645944 | 0.2903991063384 0393 | 72 | metastasis |
| cg21614638 | 1 | 7.0 | 0.016800320779998 18 | 0.2 | Metilaçk | 0.2903535948806089 | 0.5511786496020297 | 73 | metastasis |
| cg09904573 | 1 | 10.0 | 0.017082308411452475 | 0.2 | Metilaçk | 0.28924547504 98344 | 0.5702130977438842 | 74 | metastasis |
| cg12205591 | 2 | 49.0 | 0.009763924086855337 | 0.4 | Metilaçk | 0.2883595844467583 | 0.2811986148 2252767 | 75 | metastasis |
| cg09404633 | 2 | 44.0 | 0.009702109785937149 | 0.4 | Metilaçk | 0.28807168576393066 | 0.2787574710649 54 | 76 | metastasis |
| cg13558495 | 2 | 41.0 | 0.009498624442818229 | 0.4 | Metilaçk | 0.28597835941357796 | 0.27072151550557 74 | 77 | metastasis |
| cg11099899 | 2 | 42.0 | 0.009472595 58879908 | 0.4 | Metilaçk | 0.2855592414729606 | 0.2696935958075 4297 | 78 | metastasis |
| cg06066303 | 1 | 8.0 | 0.016365334519391986 | 0.2 | Metilaçk | 0.28479182423383337 | 0.5418886733720371 | 79 | metastasis |
| cg24362401 | 2 | 41.0 | 0.009394124820570944 | 0.4 | Metilaçk | 0.2847403031880 0486 | 0.266594661416333 | 80 | metastasis |
| cg27661264 | 1 | 10.0 | 0.016672488803225822 | 0.2 | Metilaçk | 0.28439042161278705 | 0.5540286781032296 | 81 | metastasis |
| cg14530601 | 2 | 44.5 | 0.009372175486162773 | 0.4 | Metilaçk | 0.28411395872127193 | 0.2657278477522 2513 | 82 | metastasis |
| cg19433435 | 2 | 47.0 | 0.009232814693032977 | 0.4 | Metilaçk | 0.2822339480 6028124 | 0.2602242713120486 | 83 | metastasis |
| cg00585790 | 2 | 46.5 | 0.009144809292689515 | 0.4 | Metilaçk | 0.28123516829557227 | 0.256748799626093 | 84 | metastasis |
| cg27105123 | 2 | 53.5 | 0.009113389851 06016 | 0.4 | Metilaçk | 0.28032212383 20906 | 0.2555079968714 6456 | 85 | metastasis |
| cg14545899 | 2 | 50.0 | 0.008990280491257139 | 0.4 | Metilaçk | 0.279115196189 7846 | 0.2506454252077787 | 86 | metastasis |
| cg14011734 | 2 | 51.0 | 0.008744832414313713 | 0.4 | Metilaçk | 0.2761320794249 77 | 0.24095308526274367 | 87 | metastasis |
| cg19570798 | 2 | 56.0 | 0.008633957996971782 | 0.4 | Metilaçk | 0.27448111608291703 | 0.2365744805103081 | 88 | metastasis |
| cg05335315 | 1 | 9.0 | 0.015558457106568613 | 0.2 | Metilaçk | 0.2730101452 9631474 | 0.5100338179877159 | 89 | metastasis |
| cg04464446 | 2 | 59.5 | 0.008410864854735243 | 0.4 | Metilaçk | 0.27163503978 10752 | 0.2277641821903 8088 | 90 | metastasis |
| cg20544605 | 2 | 61.0 | 0.008409313917894429 | 0.4 | Metilaçk | 0.2715368864297338 | 0.2277029332604 0278 | 91 | metastasis |
| cg09307264 | 1 | 10.0 | 0.015514924687 12127 | 0.2 | Metilaçk | 0.2705 782145083832 | 0.5083146544218833 | 92 | metastasis |
| cg09793001 | 2 | 58.5 | 0.008310087747299065 | 0.4 | Metilaçk | 0.2704968452 6380613 | 0.2237843605330342 | 93 | metastasis |
| cg22598744 | 2 | 55.0 | 0.00828170 8920 0503 | 0.4 | Metilaçk | 0.2703705122009 1096 | 0.2226636120982746 | 94 | metastasis |
| cg08347423 | 2 | 60.5 | 0.008300064603955687 | 0.4 | Metilaçk | 0.2702685844378873 | 0.2233885063918738 | 95 | metastasis |
| cg16510657 | 2 | 58.0 | 0.008224395754763478 | 0.4 | Metilaçk | 0.2695098979421 6546 | 0.2204032247789 6964 | 96 | metastasis |
| cg11648289 | 1 | 12.0 | 0.015623583713663954 | 0.2 | Metilaçk | 0.2691601094096137 | 0.5126049880033278 | 97 | metastasis |
| cg18416801 | 1 | 16.0 | 0.015878740614691824 | 0.2 | Metilaçk | 0.2685694007 9987107 | 0.5226823163917271 | 98 | metastasis |
| cg13983578 | 2 | 58.0 | 0.008133293108204376 | 0.4 | Metilaçk | 0.2684305619941272 | 0.2168024382855086 | 99 | metastasis |
| cg18081313 | 2 | 62.0 | 0.008144965 73080721 | 0.4 | Metilaçk | 0.2683536257161029 | 0.2172634084716 8561 | 100 | metastasis |
| cg14296147 | 2 | 58.0 | 0.008100199809 52645 | 0.4 | Metilaçk | 0.2680384901 03229 | 0.2154955319825 1473 | 101 | metastasis |
| cg20469837 | 1 | 16.0 | 0.015372090749260055 | 0.2 | Metilaçk | 0.262566881295 9561 | 0.5026739180453439 | 102 | metastasis |
| cg07075930 | 2 | 61.5 | 0.007851786030729604 | 0.4 | Metilaçk | 0.2625360904287516 | 0.1977869680958 3873 | 103 | metastasis |
| cg10864501 | 1 | 14.0 | 0.015231783637268714 | 0.2 | Metilaçk | 0.2624732243 2977144 | 0.497132969988127 | 104 | metastasis |
| cg20573242 | 2 | 63.5 | 0.007463762537828024 | 0.4 | Metilaçk | 0.260209262774231 6 | 0.19036182526811 07 | 105 | metastasis |
| cg06853304 | 2 | 60.5 | 0.007413215935157421 | 0.4 | Metilaçk | 0.2597616706 9984446 | 0.1883654605983 9762 | 106 | metastasis |
| cg06324671 | 2 | 66.5 | 0.007302522121773342 | 0.4 | Metilaçk | 0.258161159416 87473 | 0.1839939881797059 | 107 | metastasis |
| cg21929875 | 1 | 16.0 | 0.014974192354327549 | 0.2 | Metilaçk | 0.25785279166 897056 | 0.4869032859487254 | 108 | metastasis |
| cg21367040 | 1 | 18.0 | 0.014898645229 90315 | 0.2 | Metilaçk | 0.2557193592208565 | 0.48397681143794274 | 109 | metastasis |
| cg08679985 | 2 | 69.5 | 0.007091346197587547 | 0.4 | Metilaçk | 0.2553317528359 72 | 0.1756543195033436 | 110 | metastasis |
| cg03453449 | 2 | 74.5 | 0.007014311865530 57 | 0.4 | Metilaçk | 0.25443264047144 02 | 0.1726121128297 4556 | 111 | metastasis |
| cg17354052 | 2 | 72.5 | 0.006851982371420991 | 0.4 | Metilaçk | 0.2525815263372847 | 0.16620146633970176 | 112 | metastasis |
| cg03242880 | 1 | 14.0 | 0.014341485468106992 | 0.2 | Metilaçk | 0.2519254427367796 | 0.4619730980 1148747 | 113 | metastasis |
| cg11957880 | 2 | 71.5 | 0.006751296191470985 | 0.4 | Metilaçk | 0.2514261840 1118864 | 0.1622252110717792 | 114 | metastasis |
| cg24010336 | 2 | 71.0 | 0.006650524579 69928 | 0.4 | Metilaçk | 0.25025145236791885 | 0.15824558190713 69 | 115 | metastasis |
| cg18604888 | 1 | 16.0 | 0.014281011292705491 | 0.2 | Metilaçk | 0.2496474577390 0093 | 0.4596091728554 9335 | 116 | metastasis |
| cg25300386 | 2 | 73.0 | 0.006485023772863103 | 0.4 | Metilaçk | 0.2482158113107 9578 | 0.1517090953603102 | 117 | metastasis |
| cg19007602 | 2 | 73.5 | 0.006395353658998333 | 0.4 | Metilaçk | 0.2471351083 0775934 | 0.1481694918312 3352 | 118 | metastasis |
| cg12091321 | 2 | 79.0 | 0.006291773574647978 | 0.4 | Metilaçk | 0.2457233825 2101283 | 0.1440779417367 0946 | 119 | metastasis |
| cg14407437 | 1 | 20.0 | 0.014084809591432 11 | 0.2 | Metilaçk | 0.2450758698 1272686 | 0.4518395342963905 | 120 | metastasis |
| cg24567948 | 1 | 21.0 | 0.014079589025589845 | 0.2 | Metilaçk | 0.24457997108 06046 | 0.4516332066323184 | 121 | metastasis |
| cg25327247 | 2 | 79.5 | 0.006171012866 36111 | 0.4 | Metilaçk | 0.24427714580015706 | 0.1393089118348 9202 | 122 | metastasis |
| cg24995240 | 2 | 81.0 | 0.006034108196003026 | 0.4 | Metilaçk | 0.2426097238527467 | 0.13390233154167594 | 123 | metastasis |
| cg07221454 | 2 | 87.0 | 0.005895215670287947 | 0.4 | Metilaçk | 0.2407979015 8771583 | 0.1284172477166 2788 | 124 | metastasis |
| cg06324048 | 1 | 19.0 | 0.013546965 585749351 | 0.2 | Metilaçk | 0.23917905826 92686 | 0.4305968608 9975613 | 125 | metastasis |
| cg20176648 | 2 | 86.0 | 0.005836572396825851 | 0.4 | Metilaçk | 0.2377597562508 6582 | 0.1182030189205 1054 | 126 | metastasis |
| cg04001333 | 2 | 74.0 | 0.005578677627172377 | 0.4 | Metilaçk | 0.237417970806222 | 0.1158376804651844 | 127 | metastasis |
| cg00633969 | 1 | 19.0 | 0.013289233824493362 | 0.2 | Metilaçk | 0.23612558874536863 | 0.4204186291512287 | 128 | metastasis |
| cg08338368 | 2 | 84.0 | 0.005268195651195235 | 0.4 | Metilaçk | 0.23344951410272335 | 0.1036552430875 0923 | 129 | metastasis |
| cg08351105 | 2 | 87.5 | 0.005139604736805402 | 0.4 | Metilaçk | 0.2318329828241 1044 | 0.0985769860 6153568 | 130 | metastasis |
| cg09624469 | 2 | 88.5 | 0.005100139872010734 | 0.4 | Metilaçk | 0.2313401738553339 | 0.0970184568 1275167 | 131 | metastasis |
| cg26090652 | 1 | 22.0 | 0.012908093672882577 | 0.2 | Metilaçk | 0.2303058940 8928464 | 0.4053668063 8457195 | 132 | metastasis |
| cg03073814 | 2 | 87.5 | 0.005210358042578193 | 0.4 | Metilaçk | 0.2303017363 8382131 | 0.0934728312 6057196 | 133 | metastasis |
| cg15944762 | 2 | 90.0 | 0.004846881449546857 | 0.4 | Metilaçk | 0.2283028674 0794106 | 0.087016884 03379619 | 134 | metastasis |
| cg25484373 | 1 | 23.0 | 0.012710854510111393 | 0.2 | Metilaçk | 0.2276065900 2038766 | 0.3975775222 9018106 | 135 | metastasis |
| cg25842356 | 2 | 89.5 | 0.004764095168364836 | 0.4 | Metilaçk | 0.227334201909 7343 | 0.0837475238 6117687 | 136 | metastasis |
| cg15417244 | 1 | 23.0 | 0.012682515911261303 | 0.2 | Metilaçk | 0.2272708492970257 | 0.3964583865 4584125 | 137 | metastasis |
| cg05724065 | 1 | 23.0 | 0.012608553301783 68 | 0.2 | Metilaçk | 0.22638457943840938 | 0.3935374870 1692015 | 138 | metastasis |
| cg20981615 | 1 | 25.0 | 0.012618344512682038 | 0.2 | Metilaçk | 0.2258835548 8293473 | 0.3939241573 0209007 | 139 | metastasis |
| cg12901064 | 1 | 26.0 | 0.012510174876 84909 | 0.2 | Metilaçk | 0.22430311798 99751 | 0.3896523686 0855895 | 140 | metastasis |
| cg17775713 | 2 | 94.0 | 0.004506194833016026 | 0.4 | Metilaçk | 0.2241740536 1504147 | 0.0735826348 5715572 | 141 | metastasis |
| cg17229388 | 2 | 94.5 | 0.004497930895401392 | 0.4 | Metilaçk | 0.224065124535695 | 0.073236278 9933289 | 142 | metastasis |
| cg18383069 | 2 | 95.0 | 0.004302294014726847 | 0.4 | Metilaçk | 0.2217364147 9270267 | 0.0655102715 3123115 | 143 | metastasis |
| cg16746462 | 1 | 23.0 | 0.012193576921375437 | 0.2 | Metilaçk | 0.221478158874 5911 | 0.3771494184 7085925 | 144 | metastasis |
| cg19587887 | 1 | 27.0 | 0.012258506590898175 | 0.2 | Metilaçk | 0.2210568935 0500676 | 0.3797135930 0240348 | 145 | metastasis |
| cg05373692 | 1 | 24.0 | 0.011875046119747434 | 0.2 | Metilaçk | 0.2173710411591375 | 0.3645701371 9712494 | 146 | metastasis |
| cg13801416 | 1 | 26.0 | 0.011916251986477246 | 0.2 | Metilaçk | 0.2172688336 3387982 | 0.3661974207 5490797 | 147 | metastasis |
| cg21468617 | 1 | 32.0 | 0.011870202046407515 | 0.2 | Metilaçk | 0.21537425720212958 | 0.3643788371 384116 | 148 | metastasis |
| cg16220183 | 1 | 26.0 | 0.011544945 6248567 | 0.2 | Metilaçk | 0.2128640383396821 | 0.3515221031075822 | 149 | metastasis |
| cg11646704 | 2 | 98.5 | 0.003376685486874405 | 0.4 | Metilaçk | 0.2106970116 7289845 | 0.0289565380 72140558 | 150 | metastasis |
| cg15641998 | 1 | 28.0 | 0.011237012106475265 | 0.2 | Metilaçk | 0.2087085035 1972936 | 0.3393731726 5197137 | 151 | metastasis |
| cg12873941 | 1 | 29.0 | 0.011223046512508966 | 0.2 | Metilaçk | 0.2083131614 950347 | 0.3388216494 2789345 | 152 | metastasis |
| cg23900225 | 1 | 29.0 | 0.011155260634434118 | 0.2 | Metilaçk | 0.2075100702 8158786 | 0.33614467871 6404 | 153 | metastasis |
| cg13625403 | 1 | 36.0 | 0.010971294688948271 | 0.2 | Metilaçk | 0.2040692778 4657495 | 0.3288795748 0423186 | 154 | metastasis |
| cg00567749 | 1 | 32.0 | 0.010851937462519303 | 0.2 | Metilaçk | 0.203310397242 8931 | 0.3241659706 0076235 | 155 | metastasis |
| cg16404106 | 1 | 37.0 | 0.010883474570352 48 | 0.2 | Metilaçk | 0.2028865840 4227625 | 0.325411420491798 | 156 | metastasis |
| cg20603888 | 1 | 39.0 | 0.010752647529994294 | 0.2 | Metilaçk | 0.2010734566 0255543 | 0.3202448553 418514 | 157 | metastasis |
| cg13114389 | 1 | 35.0 | 0.010274486711326372 | 0.2 | Metilaçk | 0.1959640158 0482828 | 0.3013615341 6424233 | 158 | metastasis |
| cg26810336 | 1 | 35.0 | 0.010232846036601868 | 0.2 | Metilaçk | 0.1954708791 2550828 | 0.2997170785 665091 | 159 | metastasis |
| cg01703884 | 1 | 41.0 | 0.010055898161003005 | 0.2 | Metilaçk | 0.1925806435 3650511 | 0.2927291292 486678 | 160 | metastasis |
| cg07965714 | 1 | 37.0 | 0.009936503110038589 | 0.2 | Metilaçk | 0.1916873672 8527414 | 0.288014031301791 | 161 | metastasis |
| cg09120035 | 1 | 44.0 | 0.009898953879956523 | 0.2 | Metilaçk | 0.1904037904267333 | 0.2865311532742962 | 162 | metastasis |
| cg15452573 | 1 | 38.0 | 0.009821314879333916 | 0.2 | Metilaçk | 0.190167725 3150119 | 0.2834650672893559 | 163 | metastasis |
| cg16158435 | 1 | 40.0 | 0.009718077095959426 | 0.2 | Metilaçk | 0.18889446 23397655 | 0.2792880451 9759225 | 164 | metastasis |
| cg14896516 | 1 | 40.0 | 0.009658764479231594 | 0.2 | Metilaçk | 0.1879917578 4297528 | 0.2770458968 749583 | 165 | metastasis |
| cg18396533 | 1 | 36.0 | 0.009586945300784776 | 0.2 | Metilaçk | 0.1876670540 9881596 | 0.2742054958 4470187 | 166 | metastasis |
| cg01651364 | 1 | 41.0 | 0.009495787555299787 | 0.2 | Metilaçk | 0.185944749471443 | 0.2706094823 6512744 | 167 | metastasis |
| cg02322989 | 1 | 43.0 | 0.009224128849848526 | 0.2 | Metilaçk | 0.1825098306652699 | 0.2598812537327178 | 168 | metastasis |
| cg19526600 | 1 | 41.0 | 0.009185817537626 26 | 0.2 | Metilaçk | 0.1822723887 2754228 | 0.2583682798 8545835 | 169 | metastasis |
| cg15329866 | 1 | 48.0 | 0.009221160904 67449 | 0.2 | Metilaçk | 0.1820138461 2188081 | 0.2597640448 960653 | 170 | metastasis |
| cg18958531 | 1 | 45.0 | 0.009158206804266175 | 0.2 | Metilaçk | 0.1815074977 527276 | 0.2571989055 5257025 | 171 | metastasis |
| cg02774486 | 1 | 42.0 | 0.009098582267503834 | 0.2 | Metilaçk | 0.1811281294 3783574 | 0.2549232221 5712687 | 172 | metastasis |
| cg17414431 | 1 | 46.0 | 0.009044340268083589 | 0.2 | Metilaçk | 0.1800896552 8994744 | 0.2527811204 7003753 | 173 | metastasis |
| cg11050116 | 1 | 47.0 | 0.008950547782041036 | 0.2 | Metilaçk | 0.1788897988 2786127 | 0.2490771037 2039196 | 174 | metastasis |
| cg02409351 | 1 | 50.0 | 0.008955891967727 21 | 0.2 | Metilaçk | 0.178708016 1096451 | 0.2492881582 7398046 | 175 | metastasis |
| cg25465634 | 1 | 48.0 | 0.008891098296221556 | 0.2 | Metilaçk | 0.1781004390 0184364 | 0.2467293544 9594136 | 176 | metastasis |
| cg16660717 | 1 | 49.0 | 0.008871128853586 97 | 0.2 | Metilaçk | 0.17778221 89595883 | 0.2459407298 652943 | 177 | metastasis |
| cg24879335 | 1 | 52.0 | 0.008875268269810 77 | 0.2 | Metilaçk | 0.1777500783 7678053 | 0.2460850115 7040054 | 178 | metastasis |
| cg20083871 | 1 | 48.0 | 0.008793727031975 74 | 0.2 | Metilaçk | 0.1769468357 8859047 | 0.2428840104 5176414 | 179 | metastasis |
| cg04755662 | 1 | 46.0 | 0.008778736371737272 | 0.2 | Metilaçk | 0.1769429208 7450297 | 0.2422920057 5188933 | 180 | metastasis |
| cg02053171 | 1 | 50.0 | 0.008793670570263935 | 0.2 | Metilaçk | 0.1767861028 3449636 | 0.2428817809 9015126 | 181 | metastasis |
| cg07816439 | 1 | 47.0 | 0.008768809742246126 | 0.2 | Metilaçk | 0.1767342934 4468716 | 0.2418920892 6006828 | 182 | metastasis |
| cg12472775 | 1 | 54.0 | 0.008808614161067 03 | 0.2 | Metilaçk | 0.1766779415 996087 | 0.2434719265 4415023 | 183 | metastasis |
| cg26991199 | 1 | 51.0 | 0.008763575912394225 | 0.2 | Metilaçk | 0.1763541424 7181902 | 0.2416932954 1888367 | 184 | metastasis |
| cg00269115 | 1 | 49.0 | 0.008725930930419568 | 0.2 | Metilaçk | 0.176061990 7996619 | 0.2402068 35998873 | 185 | metastasis |
| cg13759778 | 1 | 53.0 | 0.008715049656480818 | 0.2 | Metilaçk | 0.1750367789 2773605 | 0.2397769174 134411 | 186 | metastasis |
| cg22813229 | 1 | 52.0 | 0.008703319558766198 | 0.2 | Metilaçk | 0.1755676881 3913524 | 0.2393136774 4491547 | 187 | metastasis |
| cg15223781 | 1 | 50.0 | 0.008653912342732223 | 0.2 | Metilaçk | 0.1751303213 0995815 | 0.2373625089 416906 | 188 | metastasis |
| cg14807303 | 1 | 54.0 | 0.008539904931591964 | 0.2 | Metilaçk | 0.1734944168404903 | 0.2328601773 4708886 | 189 | metastasis |
| cg06806711 | 1 | 55.0 | 0.008522414412010808 | 0.2 | Metilaçk | 0.1732226335494037 | 0.2321694492 7837269 | 190 | metastasis |
| cg04118154 | 1 | 52.0 | 0.008500950605746531 | 0.2 | Metilaçk | 0.1731701278 7046265 | 0.2313218098 826743 | 191 | metastasis |
| cg09088834 | 1 | 49.0 | 0.008481533445193186 | 0.2 | Metilaçk | 0.1731664987 2146123 | 0.2305549957 382041 | 192 | metastasis |
| cg04893087 | 1 | 53.0 | 0.008388021951456964 | 0.2 | Metilaçk | 0.1717823277 3045136 | 0.2268620800 8915887 | 193 | metastasis |
| cg22774472 | 1 | 57.0 | 0.008405259125595803 | 0.2 | Metilaçk | 0.171711116 8066926 | 0.2275428033 14874548 | 194 | metastasis |
| cg10983208 | 1 | 50.0 | 0.008318679580613 27 | 0.2 | Metilaçk | 0.1711588609 3804538 | 0.2241236410 3531465 | 195 | metastasis |
| cg00712898 | 1 | 55.0 | 0.008323329842516786 | 0.2 | Metilaçk | 0.1708636147 2429681 | 0.2243072871 7622753 | 196 | metastasis |
| cg19147390 | 1 | 60.0 | 0.008315262916818937 | 0.2 | Metilaçk | 0.1704753020 1053052 | 0.2239887116 1980122 | 197 | metastasis |
| cg14396117 | 1 | 58.0 | 0.008287891440519 22 | 0.2 | Metilaçk | 0.1702621612 272618 | 0.2229077690 6262397 | 198 | metastasis |
| cg20580069 | 1 | 52.0 | 0.008254391034082128 | 0.2 | Metilaçk | 0.1702490205 3624624 | 0.2215847854 3528626 | 199 | metastasis |
| cg20103758 | 1 | 59.0 | 0.008273684112284622 | 0.2 | Metilaçk | 0.1700373431 4522753 | 0.2223468993 7298063 | 200 | metastasis |
| cg18346038 | 1 | 59.0 | 0.008235992580611346 | 0.2 | Metilaçk | 0.1695907938 2302397 | 0.2208582016 3230216 | 201 | metastasis |
| cg10245048 | 1 | 55.0 | 0.008201592625768004 | 0.2 | Metilaçk | 0.1694213365 9609065 | 0.2194989934 265403 | 202 | metastasis |
| cg01473816 | 1 | 60.0 | 0.008187012734654398 | 0.2 | Metilaçk | 0.1689626970 2003 4054 | 0.2189476055 858346 | 203 | metastasis |
| cg16923271 | 1 | 60.0 | 0.008183454678178057 | 0.2 | Metilaçk | 0.1689137077 475096 | 0.2187833974 0973154 | 204 | metastasis |
| cg00342530 | 1 | 61.0 | 0.008178307537828961 | 0.2 | Metilaçk | 0.1687998450 789968 | 0.2185801287 512797 | 205 | metastasis |
| cg14448104 | 1 | 62.0 | 0.008175313917736645 | 0.2 | Metilaçk | 0.1687131749 7502383 | 0.2184619060 0340218 | 206 | metastasis |
| cg14013604 | 1 | 61.0 | 0.008105508626408932 | 0.2 | Metilaçk | 0.1686482103 4960648 | 0.2180746796 5997854 | 207 | metastasis |
| cg15071391 | 1 | 55.0 | 0.008028263051869343 | 0.2 | Metilaçk | 0.1673678195 2792487 | 0.2126546365 2165437 | 208 | metastasis |
| cg23202409 | 1 | 63.0 | 0.008042187092343557 | 0.2 | Metilaçk | 0.1670863558 1991045 | 0.2132045187 3303484 | 209 | metastasis |
| cg05661333 | 1 | 56.0 | 0.007977858329364675 | 0.2 | Metilaçk | 0.1667079944 0653483 | 0.2106640749 2236758 | 210 | metastasis |
| cg05417615 | 1 | 57.0 | 0.007924540913617164 | 0.2 | Metilaçk | 0.1660158218 107493 | 0.2085584864 9580108 | 211 | metastasis |
| cg26738080 | 1 | 57.0 | 0.007917389829903964 | 0.2 | Metilaçk | 0.165931099 6550234 | 0.2082760789 7651468 | 212 | metastasis |
| cg03854071 | 1 | 58.0 | 0.007903499940942284 | 0.2 | Metilaçk | 0.1657080941 4706356 | 0.2077275454 619632 | 213 | metastasis |
| cg24834740 | 1 | 64.0 | 0.007898974740547607 | 0.2 | Metilaçk | 0.1653415745 3581666 | 0.2075488381 9631192 | 214 | metastasis |
| cg10712623 | 1 | 64.0 | 0.007859628666308261 | 0.2 | Metilaçk | 0.1648754230 6953187 | 0.2059949996 7538264 | 215 | metastasis |
| cg11618577 | 1 | 61.0 | 0.007835862488185956 | 0.2 | Metilaçk | 0.1647427373 4136358 | 0.2050564365 2916892 | 216 | metastasis |
| cg20199333 | 1 | 62.0 | 0.007807943769848575 | 0.2 | Metilaçk | 0.1643807678 6312015 | 0.2039538822 9505657 | 217 | metastasis |
| cg25201363 | 1 | 63.0 | 0.007801205960598852 | 0.2 | Metilaçk | 0.1642313386 8894363 | 0.2036877956 3314549 | 218 | metastasis |
| cg13337865 | 1 | 65.0 | 0.007772127373818215 | 0.2 | Metilaçk | 0.1637921340 1145984 | 0.2025394366 0385607 | 219 | metastasis |
| cg17188046 | 1 | 64.0 | 0.007724756764525806 | 0.2 | Metilaçk | 0.1632775321 4608412 | 0.2006688968 9720346 | 220 | metastasis |
| cg25671438 | 1 | 63.0 | 0.007683582768214775 | 0.2 | Metilaçk | 0.1628378013 487337 | 0.1990426711 624457 | 221 | metastasis |
| cg17307280 | 1 | 66.0 | 0.007651348053877868 | 0.2 | Metilaçk | 0.1623159761 3764026 | 0.1977696717 0258192 | 222 | metastasis |
| cg20264534 | 1 | 59.0 | 0.007549285210099049 | 0.2 | Metilaçk | 0.1614550482 791706 | 0.1937390498 1945758 | 223 | metastasis |
| cg10599444 | 1 | 67.0 | 0.007528229653076688 | 0.2 | Metilaçk | 0.1608134363 187027 | 0.1929075328 2704828 | 224 | metastasis |
| cg16384137 | 1 | 68.0 | 0.007493142631987055 | 0.2 | Metilaçk | 0.1603551181 1372957 | 0.1915218912 9697297 | 225 | metastasis |
| cg22820108 | 1 | 64.0 | 0.007471537897752666 | 0.2 | Metilaçk | 0.1602775290 0728984 | 0.1906688864 345558 | 226 | metastasis |
| cg11847808 | 1 | 70.0 | 0.007414088424026372 | 0.2 | Metilaçk | 0.1593368763 7439817 | 0.1883999165 531582 | 227 | metastasis |
| cg15343119 | 1 | 67.0 | 0.007379269397958538 | 0.2 | Metilaçk | 0.1590486340 4071756 | 0.1870248585 6709763 | 228 | metastasis |
| cg01599709 | 1 | 71.0 | 0.007368674652051077 | 0.2 | Metilaçk | 0.1587597144 0477146 | 0.1866064554 233123 | 229 | metastasis |
| cg16986720 | 1 | 68.0 | 0.007357770065526389 | 0.2 | Metilaçk | 0.1587512955 808198 | 0.1861758161 8727372 | 230 | metastasis |
| cg08729012 | 1 | 70.0 | 0.007337064776455567 | 0.2 | Metilaçk | 0.1584243409 582501 | 0.1853581318 2599803 | 231 | metastasis |
| cg11768886 | 1 | 67.0 | 0.007312844528478224 | 0.2 | Metilaçk | 0.1582616673 3687608 | 0.1844016362 2095945 | 232 | metastasis |
| cg00579393 | 1 | 68.0 | 0.007312689820377409 | 0.2 | Metilaçk | 0.1582169717 417966 | 0.1843947367 2386306 | 233 | metastasis |
| cg16822189 | 1 | 71.0 | 0.007320685909195506 | 0.2 | Metilaçk | 0.1581911691 7341441 | 0.1847113046 5212213 | 234 | metastasis |
| cg08578641 | 1 | 71.0 | 0.007230862265163849 | 0.2 | Metilaçk | 0.157126986170517 | 0.1811640279 757975 | 235 | metastasis |
| cg16891845 | 1 | 61.0 | 0.007175399802352616 | 0.2 | Metilaçk | 0.1589179249 3379653 | 0.1789737282 739454 | 236 | metastasis |
| cg06220521 | 1 | 72.0 | 0.007197860453592149 | 0.2 | Metilaçk | 0.1586979464 2006107 | 0.1798607348 4221272 | 237 | metastasis |
| cg01297972 | 1 | 69.0 | 0.007098689269725319 | 0.2 | Metilaçk | 0.1556168590 7437914 | 0.1758857207 241209 | 238 | metastasis |
| cg11759378 | 1 | 73.0 | 0.007105399570748587 | 0.2 | Metilaçk | 0.1555654954 8256196 | 0.1762093092 6619752 | 239 | metastasis |
| cg18433694 | 1 | 74.0 | 0.007106408924512892 | 0.2 | Metilaçk | 0.1555414177 3581004 | 0.1762491702 3047767 | 240 | metastasis |
| cg26365854 | 1 | 76.0 | 0.007097658568985228 | 0.2 | Metilaçk | 0.1553684840 8464202 | 0.1759030049 5746474 | 241 | metastasis |
| cg05019152 | 1 | 64.0 | 0.007057049437852981 | 0.2 | Metilaçk | 0.1553668890 6746615 | 0.1742998860 3514354 | 242 | metastasis |
| cg05568762 | 1 | 72.0 | 0.006987206069142134 | 0.2 | Metilaçk | 0.1542022247 4416792 | 0.1715416623 8923554 | 243 | metastasis |
| cg21628553 | 1 | 74.0 | 0.006962615593852658 | 0.2 | Metilaçk | 0.1538378304 4464322 | 0.1705705459 2658843 | 244 | metastasis |
| cg15817236 | 1 | 77.0 | 0.006971136826209741 | 0.2 | Metilaçk | 0.1538362213 9390207 | 0.1709070627 6599843 | 245 | metastasis |
| cg14178895 | 1 | 80.0 | 0.006880450665198117 | 0.2 | Metilaçk | 0.1526668529 8460703 | 0.167325723 9404596 | 246 | metastasis |
| cg06938878 | 1 | 76.0 | 0.006747763542685819 | 0.2 | Metilaçk | 0.1512231129 654862 | 0.1620857012 2694535 | 247 | metastasis |
| cg03819692 | 1 | 83.0 | 0.006753940978189898 | 0.2 | Metilaçk | 0.1510798497 3600144 | 0.1623296578 5016354 | 248 | metastasis |
| cg16944093 | 1 | 79.0 | 0.006701395169586263 | 0.2 | Metilaçk | 0.1505763624 2015177 | 0.1602545414 0050582 | 249 | metastasis |
| cg26001336 | 1 | 67.0 | 0.006528970070065759 | 0.2 | Metilaçk | 0.1490932122 614093 | 0.1538401193 0273694 | 250 | metastasis |
| cg07845392 | 1 | 78.0 | 0.006529155264039228 | 0.2 | Metilaçk | 0.1486858754 406407 | 0.1538474329 034015 | 251 | metastasis |
| cg07015629 | 1 | 82.0 | 0.006547064464897517 | 0.2 | Metilaçk | 0.1486575724 5172867 | 0.1541597796 583727 | 252 | metastasis |
| cg24403722 | 1 | 83.0 | 0.006505002947754145 | 0.2 | Metilaçk | 0.1481305636 8086083 | 0.1524987043 330282 | 253 | metastasis |
| cg13344740 | 1 | 79.0 | 0.006488216186582921 | 0.2 | Metilaçk | 0.1480507306 3629276 | 0.1518357687 8764246 | 254 | metastasis |
| cg17769158 | 1 | 80.0 | 0.006457007894269841 | 0.2 | Metilaçk | 0.1477680181 437387 | 0.1509982200 3968247 | 255 | metastasis |
| cg18701598 | 1 | 81.0 | 0.006453733311825227 | 0.2 | Metilaçk | 0.1475812201 7379832 | 0.1504739859 4518133 | 256 | metastasis |
| cg16342780 | 1 | 84.0 | 0.006442526989067214 | 0.2 | Metilaçk | 0.1473623703 7798947 | 0.1500314306 7172955 | 257 | metastasis |
| cg05876083 | 1 | 69.0 | 0.006311918113823516 | 0.2 | Metilaçk | 0.1463191872 185087 | 0.1448734912 045528 | 258 | metastasis |
| cg22473973 | 1 | 85.0 | 0.006353923319977363 | 0.2 | Metilaçk | 0.1462852812 1151196 | 0.1465323327 2054378 | 259 | metastasis |
| cg21341807 | 1 | 70.0 | 0.006302516823014552 | 0.2 | Metilaçk | 0.1461675642 5115238 | 0.1445022094 7567227 | 260 | metastasis |
| cg10441365 | 1 | 79.0 | 0.006311559083543089 | 0.2 | Metilaçk | 0.1459577907 6058985 | 0.1448593025 3529944 | 261 | metastasis |
| cg16129797 | 1 | 71.0 | 0.006203408552443996 | 0.2 | Metilaçk | 0.1449542582 7545488 | 0.1405882683 255904 | 262 | metastasis |
| cg15572787 | 1 | 83.0 | 0.006192398791778966 | 0.2 | Metilaçk | 0.1444269950 6025772 | 0.1401534755 9768448 | 263 | metastasis |
| cg01278696 | 1 | 72.0 | 0.006126877520825624 | 0.2 | Metilaçk | 0.1440095073 600553 | 0.1375659377 7552686 | 264 | metastasis |
| cg23640701 | 1 | 89.0 | 0.006146655526984756 | 0.2 | Metilaçk | 0.1437263347 5348243 | 0.1383470417 7086737 | 265 | metastasis |
| cg08724474 | 1 | 73.0 | 0.005897248847171907 | 0.2 | Metilaçk | 0.1424367121 049568 | 0.1324468980 075136 | 266 | metastasis |
| cg21758545 | 1 | 81.0 | 0.005988498950394721 | 0.2 | Metilaçk | 0.1420693699 4502697 | 0.1321011508 5127686 | 267 | metastasis |
| cg12680071 | 1 | 75.0 | 0.005943872622026435 | 0.2 | Metilaçk | 0.1417332150 7896544 | 0.1303387871 0532334 | 268 | metastasis |
| cg19410364 | 1 | 91.0 | 0.005813748008651785 | 0.2 | Metilaçk | 0.1397339016 1329164 | 0.1251999618 9937896 | 269 | metastasis |
| cg11738011 | 1 | 84.0 | 0.005764031328851 68 | 0.2 | Metilaçk | 0.1393239128 323733 | 0.1232365721 8634242 | 270 | metastasis |
| cg15304928 | 1 | 85.0 | 0.005707423800477 06 | 0.2 | Metilaçk | 0.1386258970 0980333 | 0.1210010520 4827162 | 271 | metastasis |
| cg09744051 | 1 | 92.0 | 0.005688861914056329 | 0.2 | Metilaçk | 0.1382285723 467201 | 0.1202601157 0770499 | 272 | metastasis |
| cg00563229 | 1 | 78.0 | 0.005625598135017052 | 0.2 | Metilaçk | 0.1378625339 990655 | 0.1177696280 9815075 | 273 | metastasis |
| cg19120125 | 1 | 86.0 | 0.005637820037103 63 | 0.2 | Metilaçk | 0.1377745376 326685 | 0.1182522901 9318624 | 274 | metastasis |
| cg22858861 | 1 | 86.0 | 0.005623328275499177 | 0.2 | Metilaçk | 0.1376028469 0972185 | 0.1176798877 8336398 | 275 | metastasis |
| cg15785580 | 1 | 79.0 | 0.005575989787962277 | 0.2 | Metilaçk | 0.1372429180 3492134 | 0.1158097267 830711 | 276 | metastasis |
| cg10332959 | 1 | 87.0 | 0.005502896388111202 | 0.2 | Metilaçk | 0.1368607585 8444452 | 0.1152934377 0572421 | 277 | metastasis |
| cg00420715 | 1 | 80.0 | 0.005516113735924214 | 0.2 | Metilaçk | 0.1365029110 0684894 | 0.1134459173 4793265 | 278 | metastasis |
| cg05958582 | 1 | 94.0 | 0.005522383007783634 | 0.2 | Metilaçk | 0.1362133133 7778145 | 0.1136935007 3295562 | 279 | metastasis |
| cg05139924 | 1 | 83.0 | 0.005367616756286221 | 0.2 | Metilaçk | 0.1346554140 6856119 | 0.1075815389 5869597 | 280 | metastasis |
| cg04099673 | 1 | 84.0 | 0.005336002185000807 | 0.2 | Metilaçk | 0.1342528500 3436122 | 0.1063330295 2630205 | 281 | metastasis |
| cg01305625 | 1 | 96.0 | 0.005270814740342 32 | 0.2 | Metilaçk | 0.1331894581 7803842 | 0.1037586750 2845114 | 282 | metastasis |
| cg06100324 | 1 | 88.0 | 0.005228475996171687 | 0.2 | Metilaçk | 0.1328731864 8304777 | 0.1020866515 72706 | 283 | metastasis |
| cg10486998 | 1 | 90.0 | 0.005069803368080149 | 0.2 | Metilaçk | 0.1309439283 3609486 | 0.0958204204 6097554 | 284 | metastasis |
| cg13892015 | 1 | 90.0 | 0.004946259744593 75 | 0.2 | Metilaçk | 0.1294802488 6007356 | 0.0909414888 7423785 | 285 | metastasis |
| cg07684796 | 1 | 91.0 | 0.004849954557258724 | 0.2 | Metilaçk | 0.1283153868 0760418 | 0.0871382458 8041973 | 286 | metastasis |
| cg04809136 | 1 | 87.0 | 0.004827356012530306 | 0.2 | Metilaçk | 0.128146465 4372852 | 0.0862457938 8185974 | 287 | metastasis |
| cg12179044 | 1 | 89.0 | 0.004739007510525356 | 0.2 | Metilaçk | 0.1270492540 8802064 | 0.0827567728 7932809 | 288 | metastasis |
| cg16991589 | 1 | 93.0 | 0.004685922228132212 | 0.2 | Metilaçk | 0.1263257651 0338211 | 0.0806603517 6304673 | 289 | metastasis |
| cg14634336 | 1 | 90.0 | 0.004677111424705871 | 0.2 | Metilaçk | 0.1262915219 9009572 | 0.0803123993 0764503 | 290 | metastasis |
| cg13563405 | 1 | 92.0 | 0.004675479158445072 | 0.2 | Metilaçk | 0.1262249191 9978346 | 0.0802479385 5124951 | 291 | metastasis |
| cg05863612 | 1 | 94.0 | 0.004600129724527054 | 0.2 | Metilaçk | 0.1259677927 5260256 | 0.0796417653 158927 | 292 | metastasis |
| cg10073875 | 1 | 91.0 | 0.004396965024803582 | 0.2 | Metilaçk | 0.1229489619 5771472 | 0.0692501630 4745488 | 293 | metastasis |
| cg01254505 | 1 | 92.0 | 0.004396293302876683 | 0.2 | Metilaçk | 0.1229172729 1858233 | 0.0692224509 4724574 | 294 | metastasis |
| cg12738424 | 1 | 96.0 | 0.004382474827744695 | 0.2 | Metilaçk | 0.1226648769 7317582 | 0.0686767376 7690907 | 295 | metastasis |
| cg15107670 | 1 | 95.0 | 0.004367630111445435 | 0.2 | Metilaçk | 0.1225104822 9300683 | 0.0680904965 3224497 | 296 | metastasis |
| cg14717170 | 1 | 96.0 | 0.004345554731978595 | 0.2 | Metilaçk | 0.1222274672 1538666 | 0.0672187051 5094518 | 297 | metastasis |
| cg12699371 | 1 | 97.0 | 0.004334339718700667 | 0.2 | Metilaçk | 0.1220735583 2909741 | 0.0667758066 7522266 | 298 | metastasis |
| cg25358938 | 1 | 97.0 | 0.004303716365900126 | 0.2 | Metilaçk | 0.1217107490 4994392 | 0.0655664424 1137767 | 299 | metastasis |
| cg23652939 | 1 | 93.0 | 0.004155345821465278 | 0.2 | Metilaçk | 0.1200397766 0001737 | 0.0597070567 5183093 | 300 | metastasis |
| cg01255591 | 1 | 98.0 | 0.004112585091722514 | 0.2 | Metilaçk | 0.1194257125 4726968 | 0.0580183684 2355884 | 301 | metastasis |
| cg09600437 | 1 | 99.0 | 0.004048796728240693 | 0.2 | Metilaçk | 0.1186497797 511478 | 0.0554992658 37159265 | 302 | metastasis |
| cg00943950 | 1 | 97.0 | 0.003764788669 4661 | 0.2 | Metilaçk | 0.1153258188 846346 | 0.0442833418 60346624 | 303 | metastasis |
| cg11600114 | 1 | 99.0 | 0.003373659950344 95 | 0.2 | Metilaçk | 0.1106511164 6585128 | 0.0288370548 86170926 | 304 | metastasis |
| cg04818845 | 1 | 100.0 | 0.003200620249671 26 | 0.2 | Metilaçk | 0.1085812316 7874228 | 0.0220034455 29800988 | 305 | metastasis |
| cg14769804 | 1 | 100.0 | 0.002706744914108183 | 0.2 | Metilaçk | 0.1027300580 4486879 | 0.0024995334 16889343 | 306 | metastasis |
| cg25944700 | 1 | 100.0 | 0.002643452078400615 | 0.2 | Metilaçk | 0.1019801980 1980199 | 0.0 | 307 | metastasis |
